# An organoid CRISPRi screen revealed that SOX9 primes human fetal lung tip progenitors to receive WNT and RTK signals

**DOI:** 10.1101/2022.01.27.478034

**Authors:** Dawei Sun, Oriol Llora Batlle, Jelle van den Ameele, John C. Thomas, Peng He, Kyungtae Lim, Walfred Tang, Chufan Xu, Kerstin B. Meyer, Sarah A. Teichmann, John C. Marioni, Stephen P. Jackson, Andrea H. Brand, Emma L. Rawlins

## Abstract

The balance between self-renewal and differentiation in human fetal lung epithelial progenitors controls the size and function of the adult organ. Moreover, progenitor cell gene regulation networks are employed by both regenerating and malignant lung cells, where modulators of their effects could potentially be of therapeutic value. Details of the molecular networks controlling human lung progenitor self-renewal remain unknown. We performed the first CRISPRi screen in primary human lung organoids to identify transcription factors controlling progenitor self-renewal. We show that SOX9 promotes proliferation of lung progenitors and inhibits precocious airway differentiation. Moreover, by identifying direct transcriptional targets using Targeted DamID we place SOX9 at the centre of a transcriptional network which amplifies WNT and RTK signalling to stabilise the progenitor cell state. In addition, the proof-of-principle CRISPRi screen and Targeted DamID tools establish a new approach for using primary human organoids to elucidate detailed functional mechanisms underlying normal development and disease.

**Highlights:**

- A pooled CRISPRi screen in human fetal lung organoids identified transcription factors controlling progenitor cell self-renewal.
- SOX9 promotes tip progenitor cell proliferation and supresses precocious airway differentiation.
- Targeted DamID (TaDa) identified SOX9 direct binding targets, revealing that SOX9 lies at the intersection of WNT and RTK signalling.
- SOX9 and ETVs co-regulate the human fetal lung progenitor self-renewal programme.

## INTRODUCTION

Respiratory disease is a leading cause of human mortality and morbidity worldwide (Labaki and Han, 2020). Multiple studies suggest that changes in human lung development can contribute to adult-onset respiratory disease (Sakornsakolpat et al., 2019; Smith et al., 2018). Lung adenocarcinomas often take on an embryonic progenitor phenotype (Laughney et al., 2020; Pacheco-Pinedo et al., 2011). Moreover, it is speculated that a long-term cure for many respiratory diseases would be to trigger developmental processes in diseased lungs to regenerate the tissue (Kotton and Morrisey, 2014; Morrisey and Hogan, 2010; Nikolić et al., 2018). It is therefore crucial to understand the detailed molecular networks which control human fetal lung progenitor self-renewal and differentiation.

Mouse research has revealed that cells located at the distal tips of the branching lung epithelium are multipotent progenitors which give rise to all lung epithelial lineages during development (Alanis et al., 2014; Rawlins et al., 2009). The cell signalling and transcription factor (TF) networks controlling mouse tip progenitors are increasingly well described (Chang et al., 2013; Gerner-Mauro et al., 2020; Herriges et al., 2015; Okubo et al., 2005; Ostrin et al., 2018; Rockich et al., 2013; Y. Zhang et al., 2008). Tip progenitor cells have recently been reported to exist in developing human lung (Danopoulos et al., 2018; Miller et al., 2018; Nikolić et al., 2017). Moreover, induced pluripotent stem cell (iPSC)-based and human tissue derived 3D organoid culture systems have been established to recapitulate key features of the *in vivo* human tip progenitors (Chen et al., 2017; Miller et al., 2018; Nikolić et al., 2017). These *in vitro* models provide new opportunities to elucidate the details of the molecular events which control human lung development. However, accounting for the variable genetic background of human samples to reach a robust conclusion on biological questions remains challenging.

CRISPR genetic perturbation tools have transformed biology research (Cho et al., 2013; Cong et al., 2013; Jinek et al., 2012; Mali et al., 2013). Large scale functional genomic studies via genome-wide screens in cancer cell lines have revealed previously unappreciated genetic regulators and accelerated the drug discovery process (Bowden et al., 2020; Doench et al., 2016; Sanjana et al., 2014). CRISPR screens have started to be applied to 3D organoid systems which can more faithfully recapitulate *in vivo* biological processes (Murakami et al., 2021; Planas-Paz et al., 2019; Ringel et al., 2020), but so far have required large cell numbers to overcome variability. CRISPR interference (CRISPRi) is particularly useful for the generation of homogenous knockdowns to study essential gene function (Gilbert et al., 2014; 2013; Mandegar et al., 2016). CRISPRi screens have been established in cancer cell lines and iPSCs (Gilbert et al., 2014; Horlbeck et al., 2016; Tian et al., 2021; 2019), but not in organoid cultures.

We used a CRISPRi screen to probe systematically the function of 49 transcription factors (TFs) in human fetal lung tip progenitors. Our screen identified TFs that positively and negatively regulate tip progenitor self-renewal, including SOX9. We show that SOX9 functions by both promoting proliferation and inhibiting differentiation. Moreover, by combining an inducible CRISPRi system with Targeted DamID (TaDa) (Cheetham et al., 2018; Marshall and Brand, 2015; Southall et al., 2013), we have identified direct SOX9 transcriptional targets. These include two receptor tyrosine kinase (RTK) signalling effectors, *ETV4* and *ETV5*, which we show work with SOX9 to co-regulate the tip progenitor programme, and LGR5, which enhances WNT signalling activity. These data place SOX9 at the intersection of WNT and RTK signalling in the developing human lungs. Our use of a state-of-the-art CRISPRi screen and TaDa to study TF function in a human tissue derived organoid system will facilitate future functional studies of the coding and non-coding genome in organoid-based research.

## RESULTS

### A CRISPRi screen identified TFs that regulate lung progenitor self-renewal

To investigate systematically the TF network that regulates developing human lung epithelial tip progenitor cells, we performed a pooled CRISPRi screen in primary organoids. We used a tip progenitor cell organoid system which faithfully recapitulates key makers and signalling pathways of the *in vivo* cells (Nikolic et al., 2017). Our knockdown system (Sun et al., 2021) uses sequential lentiviral induction of an inducible CRISPRi vector, in which catalytically inactive Cas9 fused to the transcriptional repressor KRAB is controlled by doxycycline (Dox) and trimethoprim (TMP), and a constitutive gRNA (**Fig. 1A-B**). After 4-5 days of drug treatment, targeted gene knockdown can be achieved (**Fig. 1C**).

**Figure 1.**
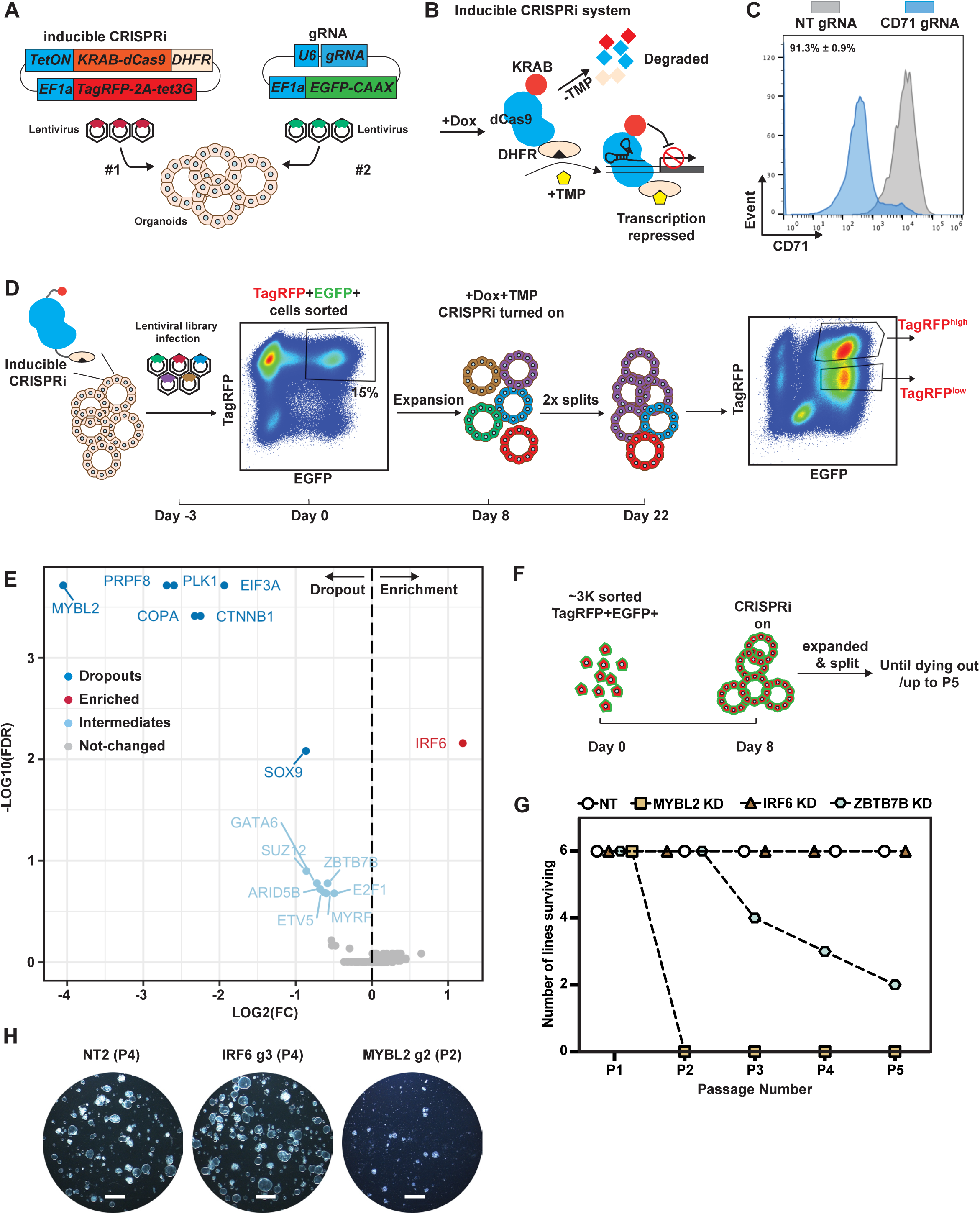
A CRISPRi screen identified crucial factors regulating human fetal lung progenitor cell self-renewal. **(A)** Schematic of introducing inducible CRISPRi system into the organoid cells using serial lentiviral infection. (**B**) Schematic of the inducible CRISPRi system. (**C**) Representative flow cytometry results showing CD71, a cell surface marker, can be efficiently knocked-down in majority of organoid cells after 5 days of Dox and TMP treatment. N = 3 different organoid lines and 3 different gRNAs for CD71 were used. Mean ± SEM is labelled. (**D**) Workflow for a focused library CRISPRi screen for transcription factors regulating human fetal lung tip progenitor cell self-renewal. Parental organoid lines with inducible CRISPRi system were established by lentiviral transduction and expanded. A CRISPRi gRNA library of 300 gRNAs was packaged into lentivirus and was used to infect single cells from two independent parental inducible CRISPRi organoid lines with an infection efficiency of ∼15%. TagRFP^+^EGFP^+^ double-positive organoid cells were collected. A fraction of cells was frozen for analysis of gRNA abundance in the starting population. The rest were seeded in Matrigel, given 8 days to recover and grown into small organoid colonies before treatment with Dox and TMP. Organoids were cultured in self-renewing medium with Dox and TMP for 2 weeks before harvest. Organoids were physically broken into pieces for passaging twice during this period. TagRFP^high^EGFP^+^ and TagRFP^low^EGFP^+^ fractions were collected separately for downstream genomic DNA isolation and analysis. (**E**) Volcano plot summarising gRNA abundance changes for each target gene. Strong depletion hits (-log10(FDR) > 2 and log2(FC) < 0) dark blue; strong enrichment hits (-log10(FDR) > 2 and log2(FC) > 0) red; intermediate depletion hits (1 < -log10(FDR) < 2 and log2(FC) < 0) light blue; unchanged genes grey. FDR, false discovery rate. (**F**) Schematic of the experimental design for serial passaging assay to validate gene knockdown effects on self-renewal. ∼3000 TagRFP^high^EGFP^+^ double positive cells were harvested for each condition and seeded in Matrigel. Organoid cells were given 8 days to recover and grown into small colonies before treating with Dox and TMP. Organoids were then maintained in Dox and TMP and serially passaged by breaking into pieces every 3-4 days. (**G**) Summary of serial passaging assay results for different knockdowns. 3 independent organoid lines each were transduced with 2 different gRNAs against the gene targets, making 6 organoid lines altogether. (**H**) Representative wide field images to show organoid growth of the indicated gene knockdown organoids at the indicated passage number. Scale bars denote 1 mm.

We designed a “drop-out” screening strategy to identify TFs which promoted, or inhibited, human fetal lung tip progenitor self-renewal (**Fig. 1D**). We curated a lentiviral CRISPRi gRNA library containing on average 5 gRNAs per gene against 49 TFs exhibiting enrichment and/or high abundance in tip progenitor cells (**Fig. S1A; Supplementary Table 1**), together with positive controls (4 essential genes) and negative controls (33 non-targeting gRNAs), 300 gRNAs total. Library sequencing confirmed that all gRNAs were successfully cloned, with the exception of one gRNA against *MYCN* (**Fig. S1B**). Two independent parental organoid lines (biological replicates) with inducible CRISPRi were dissociated to single cells and transduced with the gRNA library at ∼15% infection rate (**Fig. 1D**), to ensure that most infected cells received one gRNA. At least 500 cells were infected per gRNA. TagRFP^+^EGFP^+^ double positive cells (containing both inducible CRISPRi and gRNA) were harvested 3 days after infection and we observed highly consistent gRNA abundance between transduced parental lines (biological replicates) (**Fig. S1C**). To test the robustness of the human fetal lung organoids for such screens, we also split the TagRFP^+^EGFP^+^ cells from one of the biological replicates and assayed them separately as technical replicates. Organoid cells were recovered for 8 days, then CRISPRi was switched on for two weeks of culture. Two different cell populations (TagRFP^high^EGFP^+^ and TagRFP^low^EGFP^+^) emerged at the end of the screen (**Fig. 1D**). This was likely due to the inducible CRISPRi vector being modified by the cellular epigenetic machinery in long-term cultures. We harvested TagRFP^high/low^ populations separately for downstream gRNA abundance analysis using next-generation sequencing (NGS).

All expected gRNA sequences were present in all downstream NGS results (**Supplementary Table 2**). The consistency between the technical replicates (R = 0.79-0.89) demonstrated the feasibility of performing a pooled CRISPRi screen using ∼500 cells/gRNA in the human fetal lung organoid system (**Fig. S1D**). Our gRNA correlation dropped to 0.57 when comparing different parental lines (biological replicates) (**Fig. S1E**), as might be expected due to the variable nature of organoids from different donors. The gRNA abundance of the TagRFP^high^EGFP^+^ versus TagRFP^low^EGFP^+^ populations correlated to a similar extent between technical (0.79 and 0.72) and biological (0.58 and 0.55) replicates (**Fig. S1D-E**). This suggested that these two populations behaved similarly, despite different TagRFP levels. Therefore, we treated them as technical replicates to find the most robust changes in gRNA abundance following passaging, ‘hits’.

gRNAs targeting all 4 positive-control essential genes (*PRPF8, PLK1, COPA*, and *EIF3A*) were strongly depleted in the screen (**Fig. 1E**). Strong depletion of gRNAs targeting *MYBL2* and *CTNNB1* (**Fig. 1E**) was consistent with *MYBL2* being a central factor regulating cell proliferation and the importance of WNT signalling for tip progenitor cell growth (Hein et al., 2021; Musa et al., 2017). gRNAs targeting *SOX9* were moderately depleted suggesting a role in tip progenitor self-renewal, consistent with previous studies (Chang et al., 2013; Li et al., 2021; Rockich et al., 2013). By contrast, depletion of *SOX2* did not markedly influence progenitor self-renewal, consistent with a recent knock-out study (Sun et al., 2021). We also obtained intermediate hits, with gRNAs being mildly depleted, including *GATA6, SUZ12, ARID5B, ETV5, ZBTB7B, E2F1* and *MYRF*. This could be results from there being a few effective gRNAs buffered by the effects of non-functional gRNAs targeting the same gene; alternatively, non-specific effects of a small group of gRNAs that influenced cell viability. Perhaps surprisingly, we discovered that gRNAs targeting *IRF6* were enriched in the screen, indicating a normal function in supressing self-renewal, consistent with reports of a tumour suppressor role (Botti et al., 2011). These results indicate that the CRISPRi screen was able to effectively identify crucial factors regulating tip progenitor cell self-renewal.

### Validation of CRISPRi hits confirmed screen robustness

To validate the screen results, we cloned individual gRNAs separately into the gRNA vector and tested their effects on progenitor self-renewal (**Fig. 1F**). We selected *MYBL2, IRF6* and *ZBTB7B*/*ARID5B* as representative of strongly-depleted, enriched and intermediately-depleted genes, respectively. The gRNAs targeting *IRF6, MYBL2* and *ZBTB7B* were able to efficiently down-regulate expression of their targeted gene (**Fig. S2A**). We performed a serial passaging assay to test the effects of target gene knockdown on organoid self-renewal. *MYBL2* knockdown organoids were rapidly lost after two serial passages in all the different organoid lines tested (6/6), whereas *ZBTB7B* knockdown organoids were slowly lost in some of the organoid lines tested (4/6, by Passage 5) (**Fig. 1F-I**). These data correlated well with the relative ‘strength’ of hits in the screen. *IRF6* knockdown led to better organoid growth compared with non-targeting (NT) control organoids, consistent with its gRNA enrichment in the screen (**Fig. 1F-I**). We further tested the effects of *IRF6* and *MYBL2* knockdown on organoid proliferation using an EdU incorporation assay. *IRF6* knockdown organoids exhibited a higher proportion of EdU^+^ cells compared with NT controls (**Fig. S2B-C**). Consistent with the serial passaging data, in *MYBL2* knockdown organoids, the percentage of EdU^+^ cells was reduced (**Fig. S2B-C**). We did not observe any significant depletion of *ARID5B* mRNA following knockdown (**Fig. S2D**), suggesting that the depletion of this gRNA in the screen might be due to off-target effects. Overall, we were able to validate the strong, and some intermediate, hits in our screen confirming that the CRISPRi screening conditions were robust enough to discover TFs that regulate tip progenitor cell self-renewal.

### SOX9 promotes proliferation and suppresses differentiation to govern progenitor self-renewal

SOX9 is an established lung epithelial tip progenitor marker, but its function and downstream targets in human fetal lung progenitors have not been elucidated. We therefore used our human fetal lung progenitor organoids, which capture key TF expression patterns (**Fig. 2A**), to explore *SOX9* function. We first validated that the gRNAs used in the screen were able to knockdown *SOX9* effectively (**Fig. 2B-C**). *SOX9* knockdown organoid cells were gradually lost in a serial passaging assay in all the organoid lines tested (6/6) (**Fig. 2D-E**), confirming that SOX9 is an important regulator of tip progenitor cell self-renewal.

**Figure 2.**
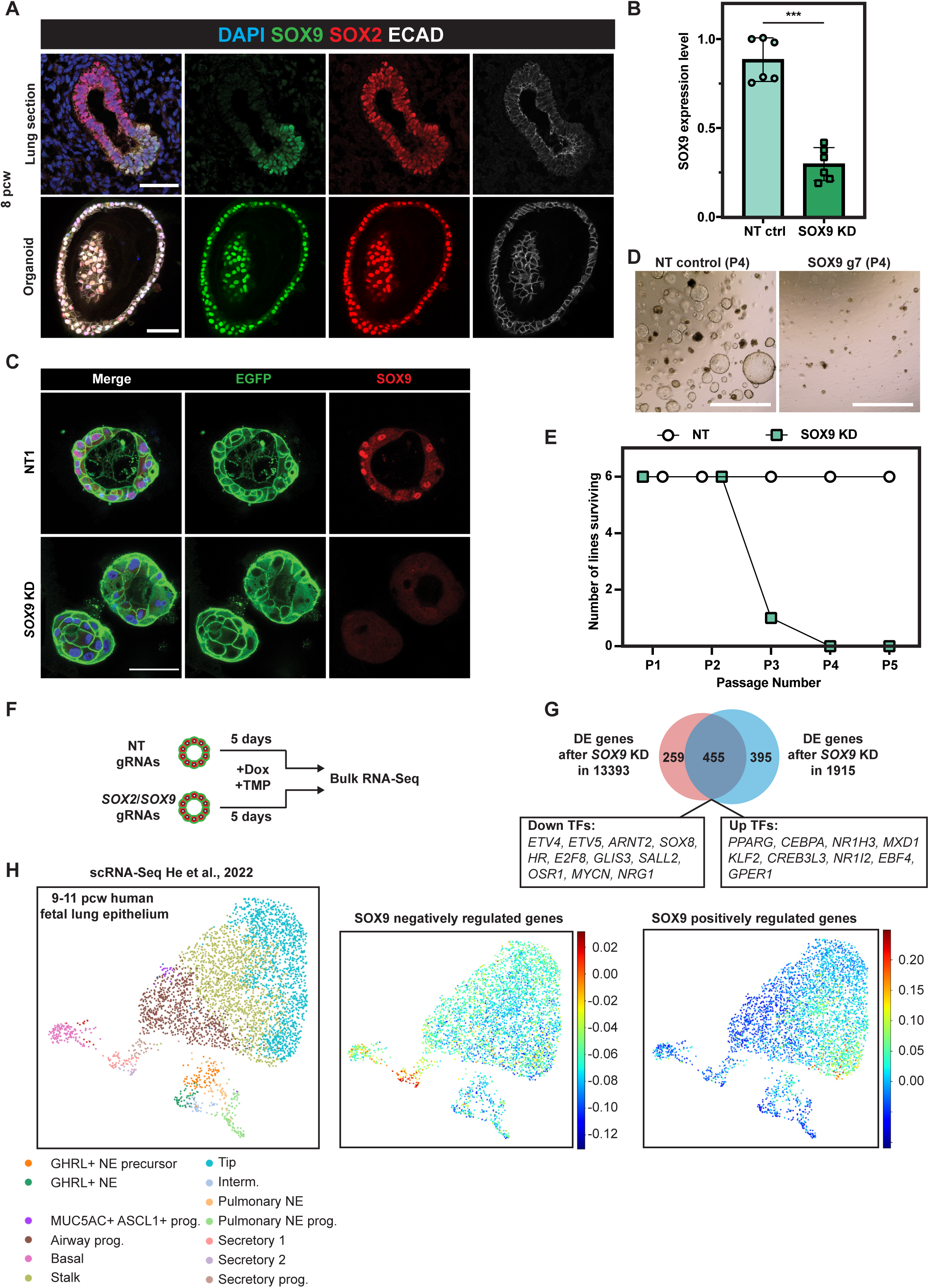
SOX9 regulates human fetal lung progenitor cell self-renewal. **(A)** Human fetal lung tip organoids faithfully express in vivo tip progenitor markers, SOX2 and SOX9. Upper panel: SOX2/SOX9 dual expression in human fetal lungs. Lower panel: SOX2/SOX9 in human fetal lung tip organoids. SOX9 green, SOX2 red, E-Cadherin white. **(B)** qRT-PCR showing that *SOX9* gRNAs effectively knocked down *SOX9* transcript levels using inducible CRISPRi after 5 days of Dox and TMP treatment. 3 different organoid lines and 2 different *SOX9* gRNAs were used. Error bars: mean ± SEM. Two-sided Student’s t-test with equal variance ***p < 0.001. **(C)** SOX9 was knocked down at the protein level using the inducible CRISPRi system after 4 days of Dox/TMP treatment. **(D)** *SOX9* knockdown organoids at Passage #4 (P4). (**E**) Summary of the serial passaging assay results for *SOX9* knockdowns. 3 independent organoid lines each were transduced with 2 different gRNAs against *SOX9*, making 6 different organoid lines altogether. Serial passaging conditions were as previously stated. (**F**) Schematic of RNA-Seq to identify SOX9 downstream targets. 2 independent inducible CRISPRi parental organoid lines were transduced with 2 different non-targeting control gRNAs, two different *SOX9* gRNAs and two different *SOX2* gRNAs respectively. Organoids were supplemented with Dox/TMP for 5 days before harvesting for RNA-Seq. (**G**) Venn Diagram showing the overlapping number of differentially expressed (DE) genes in *SOX9* knockdown organoids from different parental lines. (**H**) All SOX9 positively and negatively regulated genes were used to score against an scRNA-Seq dataset from 9-11pcw human fetal lung epithelium (left panel, He et al., 2022). SOX9 negatively-regulated genes were primarily enriched in secretory lineage populations (middle panel). SOX9 positively-regulated genes were primarily enriched in tip progenitor cells (right panel). Prog., progenitors. NE, neuroendocrine cells. Interm, intermediate NE. Scale bars denote 50 µm (**A, C**) and 100 µm (**D**).

We performed RNA-Seq on *SOX9* knockdown organoids to characterise downstream targets. To account for biological variation between organoids from different donors and identify the most robust downstream targets we used: (1) two different gRNAs targeting *SOX9* to circumvent potential gRNA-dependent non-specific effects; (2) two different NT control gRNAs to eliminate lentiviral transduction-dependent non-specific effects; (3) two independent organoid lines to avoid possible organoid line non-specific effects. *SOX2* knockdown organoids were included for comparison. Organoids were harvested after 5 days of Dox/TMP treatment to probe the early effects of *SOX9* knockdown (**Fig. 2F**).

Unsupervised hierarchical clustering of the RNA-Seq transcriptome data revealed that the two independent organoid lines clustered separately (**Fig. S3A**), indicating that there was an organoid line-dependent effect. However, within each cluster, NT controls and *SOX2* knockdown organoids clustered separately from *SOX9* knockdown organoids, suggesting that *SOX9* knockdown led to unique gene expression profiles (**Fig. S3A**). To obtain the most robust differentially expressed (DE) genes, we first identified DE genes between the *SOX9*, or *SOX2*, knockdown groups and NT controls of the same organoid line using a 2-fold change in gene expression and an adjusted P-value ≤ 0.05 as cut-offs. We then overlapped the DE gene lists between biological replicates.

For *SOX2* knockdown, this strategy identified 54 genes in organoid line 1915 and 133 genes in organoid line 13393 with an overlap of only 20 genes (including *SOX2*) (**Fig. S3B**; **Supplementary Table 3**), supporting our previous conclusion that SOX2 is dispensable for progenitor cell self-renewal (Sun et al., 2021). By contrast, 850 genes (organoid line 1915) and 714 genes (organoid line 13393) were differentially expressed after *SOX9* knockdown, with an overlap of 455 genes, including 20 TFs (**Fig. 2G; Supplementary Table 3**). For *SOX9*, more than 50% of the DE genes were shared between biological replicates (53.5% for line 1915 and 63.7% for line 13393) (**Fig. 2G**), indicating that the experimental design was able to remove most non-specific effects.

*SOX9* itself was not in the *SOX9* knockdown DE gene list (**Fig. 2G**), even though we had previously validated both mRNA and protein depletion (**Fig. 2B-C**). We hypothesised that an alternative splicing event might be enhanced by CRISPRi induced knockdown, buffering overall *SOX9* locus transcription. We therefore used sashimi plots to visualise the splice junctions (**Fig. S3C**). This showed that there was an alternative transcriptional start site(s) (TSS) within *SOX9* intron #1 and that hybrid reads of intron #1 and exon #2 emerged after *SOX9* knockdown. However, the ‘new’ *SOX9* transcript was unlikely to be functional as it lacked the dimerization domain and part of the HMG domain encoded in SOX9 exon #1 (**Fig. S3C**). Additionally, following knockdown we were unable to detect SOX9 using an antibody against the C-terminus (**Fig. 2C**).

To test whether the *SOX9* knockdown cells altered their differentiation status, we took advantage of our human fetal lung scRNA-Seq dataset (He et al., 2022). We subset the human fetal lung scRNA-Seq data from 9-11 post conception weeks (pcw) and used all SOX9 positively and negatively regulated genes to score the epithelial compartment (**Fig. 2H**). SOX9 negatively-regulated genes were primarily expressed in secretory cells and their progenitors, indicating that SOX9 suppresses premature differentiation (**Fig. 2H, middle panel**). By contrast, SOX9-positively regulated genes were primarily enriched in the tip progenitor cells, confirming that SOX9 functions to drive tip progenitor cell gene expression programmes (**Fig. 2H, right panel**).

Gene ontology (GO) revealed that genes involved in cell metabolic processes were upregulated after *SOX9* knockdown (**Fig. S3D-E; Supplementary Table 4**). Additionally, other foregut lineage markers, including from liver (*APOL1* and *ALB*) and stomach (*GKN1, GKN2* and *TFF1*), were upregulated (**Fig. S3E**). Cell division GO terms were enriched in the down-regulated genes (**Fig. S3D; Supplementary Table 5**), consistent with the observation that *SOX9* knockdown organoids were gradually lost in the serial passaging assay (**Fig. 2D-E**). Overall, our results showed that SOX9 appears to maintain tip progenitor self-renewal via two major effects: 1. promoting tip cell proliferation; 2. inhibiting differentiation by suppressing secretory cell lineage-related metabolic genes and non-lung lineage genes.

### Targeted DamID (TaDa) identified SOX9 direct transcriptional targets

To identify genes directly activated or repressed by SOX9, we used SOX9 Targeted DamID (TaDa) to probe genomic occupancy (Cheetham et al., 2018; Marshall and Brand, 2015; Marshall et al., 2016; Southall et al., 2013). We introduced the TaDa system into organoids via lentivirus and harvested cells 48-72 hours after transduction (**Fig. 3A**). 4845 SOX9 binding peaks were called with high confidence (FDR<10^−50^) versus Dam-only controls (no SOX9 fusion) and shared by 4 independent organoid lines (**Supplementary Table 6**). De novo motif analysis revealed the enrichment of a SOX motif (**Fig. S4A**), suggesting that the SOX9-Dam fusion protein faithfully bound to SOX9 targets. We overlapped the *SOX9* knockdown DE gene list with peak annotations (http://great.stanford.edu/public/html/, GREAT analysis) and identified 171 genes directly regulated by SOX9 (**Fig. 3B; Supplementary Table 6**). SOX9 positively-regulated genes enriched in tip progenitor cells and negatively-regulated genes enriched in secretory cell lineages (**Fig. S4B**). Direct transcriptional targets included *ETV4, ETV5, MYCN, LGR5, CD44, CXCR4* and *SHH*, all of which have been previously reported to be important for lung development in mouse (**Fig. 4C**) (Hein et al., 2021; Herriges et al., 2015; Okubo et al., 2005; Weaver et al., 2003). *SHH* expression in tip progenitor cells was confirmed by in situ HCR (hybridization chain reaction) (**Fig. S4C**).

**Figure 3.**
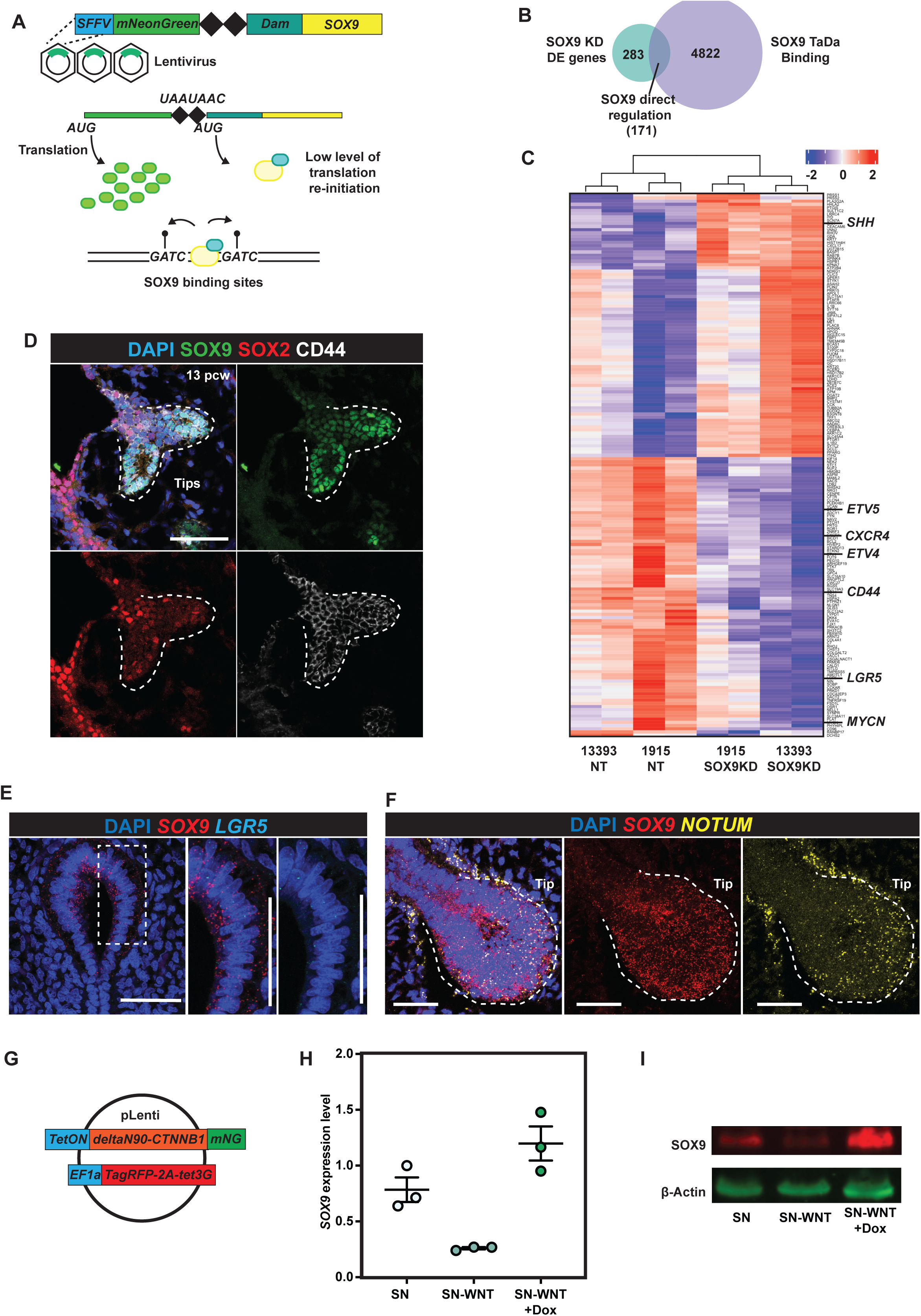
Targeted DamID (TaDa) identified SOX9 direct regulating targets. (**A**) Schematic of *SOX9* TaDa lentiviral construct design. SOX9 is fused with *E*.*coli* DNA adenine methylase (Dam). The fusion protein binds to SOX9 binding targets and methylates adenine in the sequence *GATC*. Methylated *GATC* is specifically recognised and cleaved by DpnI restriction enzyme. TaDa is designed to produce an extremely low level of Dam-Sox9 fusion protein: an mNeonGreen open reading frame (ORF1) was placed in front of *Dam-SOX9* fusion (ORF2). The two ORFs were separated by two stop codons and a frame-shift C (represented by 2 black diamonds), such that the Dam-SOX9 fusion protein is translated at very low levels after rare translational re-initiation events. (**B**) Venn Diagram showing the overlap between DE genes identified in the *SOX9* knockdown RNA-Seq experiment and genes annotated from SOX9 TaDa peaks. (**C**) Heatmap showing expression level of all 171 SOX9 directly regulated genes across non-targeting control and *SOX9* knockdown organoid lines. (**D**) CD44 protein expression in SOX9^+^ tip progenitor cells. SOX9 green, SOX2 red, CD44 white. (**E**) *LGR5* mRNA expression enrichment in *SOX9* expressing tip progenitor cells. *SOX9* red, *LGR5* cyan. (**F**) Tip progenitor cells are of high WNT signalling activity. NOTUM yellow, SOX9 red. (**G**) Design of constitutively activated β-catenin over-expression lentiviral construct. qRT-PCR showing rescue of *SOX9* transcription after constitutively activated β-catenin overexpression in organoids cultured without WNT activators. N = 3 different organoid lines. WB showing SOX9 protein was rescued after constitutively activated β-catenin overexpression in organoids cultured without WNT activators. Scale bars = 50 µm in all panels.

**Figure 4.**
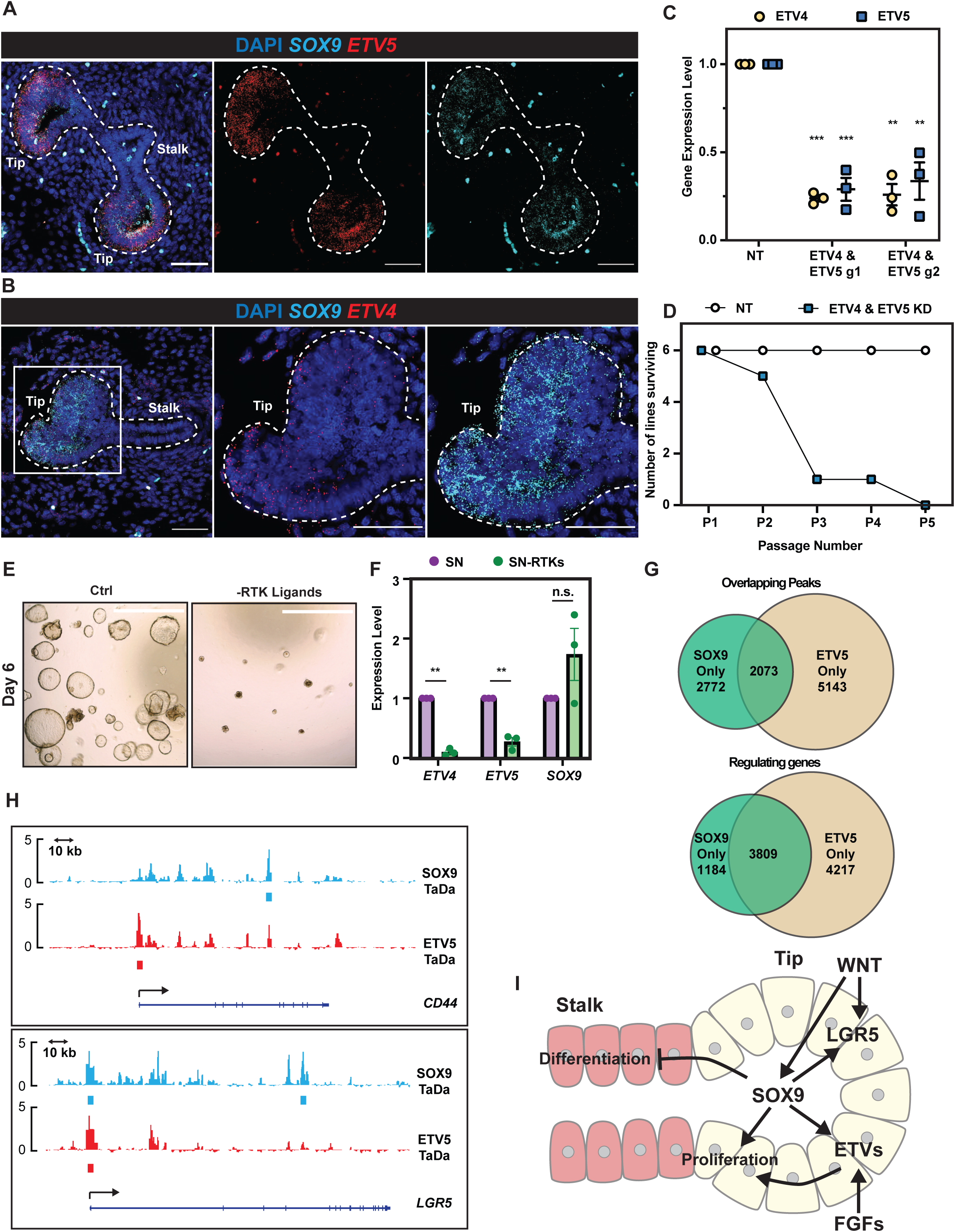
SOX9 directly regulates *ETV4* and *ETV5* expression thereby enhancing FGF signalling. (**A,B**) HCR images showing *ETV5* (A) and *ETV4* (B) expression in *SOX9* expressing tip progenitors. *ETV4/5* red, *SOX9* cyan. Scale bars = 50 µm. (**C**) qRT-PCR showing *ETV4* and *ETV5* dual knockdown in tip organoids. N =3 different organoid lines and 2 different *ETV4* and *ETV5* gRNA combinations were used. Error bars: mean ± SEM. Two tailed paired t-test: **p < 0.01; ***p < 0.001. (**D**) Summary of the serial passaging assay results for *ETV4*; *ETV5* dual knockdowns. 3 independent organoid lines each were transduced with 2 different *ETV4*; *ETV5* gRNA combinations, making 6 different organoid lines altogether. Serial passaging conditions were as previously stated. (**E**) Representative images of organoids grown in self-renewing medium (left) and self-renewing medium without RTK ligands (EGF, FGF7, FGF10) (right). (**F**) qRT-PCR of *ETV4, ETV5* and *SOX9* expression level in organoids in self-renewing medium and self-renewing medium without RTK ligands. N = 3 different organoid lines used. Error bars: mean ± SEM. Two-tailed paired T-test: **p < 0.01; n.s. non-significant. (**G**) Venn diagrams showing overlap of SOX9-ETV5 TaDa peak regions (upper) and overlapping genes from TaDa peak annotations (lower). (**H**) Examples of SOX9 and ETV5 genomic occupancies showing co-regulation. At the *CD44* locus (upper), SOX9 binds to an intron, whereas ETV5 primarily occupied the promoter. By contrast, for the *LGR5* gene (lower), SOX9 and ETV5 both showed occupancy at the promoter. (**I**) Schematic model of SOX9 regulation of tip progenitor cell self-renewal.

We confirmed CD44 protein (**Fig. 3D**) and *LGR5* mRNA localisation (**Fig. 3E**) in *SOX9*-expressing tip regions, consistent with previous reports for *LGR5* expression pattern (Hein et al., 2021; Ostrin et al., 2018). LGR5 and CD44 are WNT signalling amplifiers (Schmitt et al., 2015; Schuijers and Clevers, 2012), suggesting that SOX9 could enhance WNT signalling in tip progenitors. *CTNNB1*, encoding β-catenin, an intracellular signal transducer of the WNT signalling pathway, was a strong hit in our screen (**Fig. 1E**) and we observed that *NOTUM*, a WNT inhibitor, is enriched in tip progenitors (**Fig. 3F**) confirming that human tips are highly WNT responsive. This prompted us to investigate whether WNT signalling influenced *SOX9* expression. We removed WNT signalling activators (R-spondin1 and CHIR99021) from tip progenitor organoid culture, leading to loss of SOX9 transcript and protein (**Fig. 3H-I**). We showed that the regulation is via β-catenin, canonical WNT signalling, by overexpressing a constitutively activated β-catenin (DeltaN90-β-catenin) (Guo et al., 2012) in the absence of WNT signalling activators (**Fig. 3G**), rescuing SOX9 expression (**Fig. 3H-I**). Therefore, SOX9 is regulated by canonical WNT signalling and directly regulates *LGR5* and *CD44* to enhance WNT signalling.

We used SOX9 overexpression in tip progenitor organoids to test the predicted direct transcriptional targets (**Fig. S4D-F**). *ETV5* and *MYCN* were significantly up-regulated upon SOX9 overexpression, further supporting the direct regulation (**Fig. S4F**). However, *ETV4* and *CFTR* remained unchanged, likely due to already saturated binding at these loci (**Fig. S4F**). These results marked the first application of TaDa in organoids to identify TF binding targets.

### SOX9 and ETVs co-ordinate the tip progenitor self-renewal programme

We were intrigued to discover that *ETV4* and *ETV5* are SOX9 direct binding targets. We showed that *ETV5* and *ETV4* are enriched in *SOX9* expressing tip progenitor cells *in vivo* (**Fig. 4A-B**). *ETV4* and *ETV5* are redundant in mouse lung development (Herriges et al., 2015). This was in line with our screen results in which *ETV5* was an intermediate hit and *ETV4* gRNAs had no effect (**Fig. 1E**). To test if *ETV4* and *ETV5* function with SOX9 in human fetal lung progenitor self-renewal, we knocked down *ETV4* and *ETV5* simultaneously using inducible CRISPRi (**Fig. 4C**). *ETV4*; *ETV5* double knockdown organoids were gradually lost during serial passaging, similar to *SOX9* knockdown (**Fig. 4D**).

ETVs are RTK signalling effectors (Herriges et al., 2015; Z. Zhang et al., 2009) and RTK ligands are required for long-term tip organoid self-renewal (Nikolic et al., 2017). We reasoned that SOX9 influences RTK signalling thereby influencing progenitor self-renewal. Therefore, we tested the effects of removing RTK ligands (EGF, FGF10 and FGF7) from our self-renewal medium; both *ETV4* and *ETV5* were down-regulated and organoids could not proliferate normally (**Fig. 4E-F**). *SOX9* expression was not affected by RTK ligand removal, suggesting its transcription is not directly controlled by RTK signalling (**Fig. 4F**). These results show that SOX9 can directly activate *ETV4* and *ETV5* expression, enhancing RTK signalling.

To identify ETV4 and ETV5 binding targets, we performed TaDa. ETV4 and ETV5 TaDa exhibited >90% similarity (**Fig. S4G**), and we primarily used the ETV5 data for analysis. Motif analysis on ETV5 bound regions identified the ETS binding motif, suggesting ETV5 TaDa recapitulated ETV5 binding events (**Fig. S4H**). The SOX motif was also mildly enriched in ETV5 binding regions. We explored the possibility of SOX9 and ETV5 co-regulation of targets. Both SOX9 and ETV5 primarily bind to intronic regions (**Fig. S4I**), consistent with previous results (Shih et al., 2015). SOX9 and ETV5 binding peaks overlapped less than 50% (SOX9: 42.8%; ETV5: 28.7%). However, peak annotation by GREAT analysis revealed a much greater overlap (SOX9: 76.3%; ETV5: 47.5%) (**Fig. 4G**), suggesting SOX9 and ETV5 bind to different regions of the same loci to coordinate the tip progenitor self-renewal programme (**Fig. 4H**).

Our results suggest a model for SOX9 function in human fetal lung progenitors (**Fig. 4I**), in which it promotes self-renewal by: (1) promoting progenitor cell proliferation, via a feed-forward WNT signalling loop and by enhancing RTK signalling via ETV factors; (2) suppressing metabolic processes favourable for precocious differentiation in early lung development; (3) suppressing other non-lung lineage gene expression to maintain human fetal lung progenitor states.

## DISCUSSION

CRISPR-related screens have rarely been applied to organoids because of the difficulties manipulating 3D culture systems. We have performed the first pooled CRISPRi dropout screen in a tissue-derived organoid system, followed by detailed characterisation of selected gene function. Previous reports challenged the robustness of such screens in organoid systems (Ringel et al., 2020), or used an enormous cell number (10,000 cells per guide) (Planas-Paz et al., 2019). Here we achieved a pooled CRISPRi screen of high robustness and sensitivity with a starting population of ∼500 cells per gRNA (**Fig. 1, S1, S2**). Our ∼500 cell representation per gRNA is similar to screens carried out in cancer cell lines (Bowden et al., 2020; Joung et al., 2017). We envision that similar screens can be practically and affordably scaled-up for targeting hundreds of genes, for example the mammalian TFome (Ng et al., 2021). The CRISPRi system has been used to study non-coding genome function (Fulco et al., 2016; 2019; Gasperini et al., 2019), also enabling the use of organoid platforms for functional validation of genome-wide association studies (GWAS) of lung diseases (Wain et al., 2017).

Our study is the first to use TaDa in human organoids (**Fig. 3-4**). TaDa is independent of ChIP-grade antibodies and requires a relatively low cell number (400K cells for TaDa in this study, typically >1 million cells for ChIP-Seq) (Cheetham et al., 2018; Southall et al., 2013; Tosti et al., 2018). We discovered *ETV4* and *ETV5* (**Fig. 3C**), among others, as SOX9 directly-regulated targets. Interestingly, such regulation was not discovered in *SOX9* knockout mouse models. This could be due to gene expression profiles being masked by non-SOX9 expressing cells in the previous mouse study (Chang et al., 2013), or species-specific roles.

We have shown that WNT signalling is highly active in tip progenitor cells, that SOX9 directly regulates WNT pathway activators and that SOX9 expression is controlled by canonical WNT signalling (**Fig. 3E-H**). These findings were consistent with previous mouse research showing β-catenin knockout led to SOX9 loss in tip progenitors (Ostrin et al., 2018) and a study in which the LGR5-WNT signalling axis was blocked in human fetal lung explants reducing the number of SOX9-positive tip cells and downregulating *ETV5* (Hein et al., 2021).

We have made the first effort to systematically identify transcription factors with roles in human lung development. These approaches and results will deepen our understanding of human lung development and disease, and greatly benefit future functional organoid-based studies of the coding and non-coding genome.

## ACKNOWLEDGEMENTS

We acknowledge the Gurdon Institute Imaging Facility, Gurdon Institute NGS core and Pathology Department Flow Cytometry Facility. DS is supported by a Wellcome Trust PhD studentship (109146/Z/15/Z) and the Department of Pathology, University of Cambridge. PH holds a non-stipendiary research fellowship at St Edmund’s College, University of Cambridge. KL is supported by the Basic Science Research Program through the National Research Foundation of Korea (NRF) funded by the Ministry of Education (2018R1A6A3A03012122). ELR is supported by the MRC (MR/P009581/1; MR/S035907/1). KBM, JCM and SAT acknowledge funding from the MRC (MR/S035907/1) and from Wellcome (WT211276/Z/18/Z and Sanger core grant WT206194); Research in the Andrea Brand lab was funded by Wellcome Trust Senior Investigator Award (103792) and the Royal Society Darwin Trust Research Professorship to AHB and Wellcome Trust Postdoctoral Training Fellowship for Clinicians (105839) to JvdA. Research in the Steve Jackson lab is funded by Cancer Research UK Discovery Award DRCPGM\100005, ERC Synergy Grant 855741 (DDREAMM), and Wellcome Investigator Award [206388/Z/17/Z]. We acknowledge core funding to the Gurdon Institute from the Wellcome Trust (203144/Z/16/Z) and CRUK (C6946/A24843).

## STAR Methods

## RESOURCE AVAILABILITY

### Lead contact

Further information and requests for resources and reagents should be directed to and will be fulfilled by the lead contact, Emma L. Rawlins.

### Materials availability

Plasmids generated in this study will be deposited to Addgene. Human lung organoid lines used in this study are available from the lead contact, Emma L. Rawlins, with a completed Materials Transfer Agreement.

### Data and code availability

*SOX9* and *SOX2* CRISPRi knockdown RNA-Seq data and SOX9/ETV4/ETV5 targeted DamID data have been deposited to GEO (GSE189885) and are publicly available as of the date of publication. Any additional information required to reanalyze the data reported in this paper is available from the lead contact upon request.

## METHOD DETAILS

### Human embryonic and foetal lung tissue

The sources of human foetal lung tissues were the MRC/Wellcome Trust Human Developmental Biology Resource (www.hdbr.org; London site REC reference: 18/LO/0822; Newcastle site REC reference: 18/NE/0290; Project 200454;) and Cambridge University Hospitals NHS Foundation Trust under NHS Research Ethical Committee permission (96/085). None of the samples used for the current study had any known genetic abnormalities.

### Derivation and maintenance of human fetal lung organoid culture

Human fetal lung organoids were derived and maintained as previously reported (Nikolić et al., 2017). Briefly, human fetal lung tissues were dissociated with Dispase (8 U/ml Thermo Fisher Scientific, 17105041) at room temperature (RT) for 2 min. Mesenchyme was carefully removed by needles. Branching epithelial tips were micro-dissected, transferred into 50 µl of Matrigel (356231, Corning) and seeded in 24 well low-attachment plate wells (M9312-100EA, Greiner). The plate was incubated at 37°C for 5 min to solidify the Matrigel. 600 µl of self-renewing medium containing: N2 (1: 100, ThermoFisher Scientific, 17502–048), B27 (1: 50, ThermoFisher Scientific, 12587–010), N-acetylcysteine (1.25 mM, Merck, A9165), EGF (50 ng/ml, PeproTech, AF-100-15), FGF10 (100 ng/ml, PeproTech, 100-26), FGF7 (100 ng/ml, PeproTech, 100-19), Noggin (100 ng/ml, PeproTech, 120-10C), R-spondin (5% v/v, Stem Cell Institute, University of Cambridge), CHIR99021 (3 µM, Stem Cell Institute, University of Cambridge) and SB 431542 (10 µM, bio-techne, 1614), was added. The plate was incubated under standard tissue culture conditions (37°C, 5% CO_2_). Once formed, organoids were maintained in self-renewing medium and passaged for routine maintenance by mechanically breaking using P200 pipettes every 3-7 days.

### Whole mount immunostaining for human foetal lung organoids

Organoids were fixed with 4% paraformaldehyde (PFA) in the culture plates on ice for 30 min. After two PBS washes, 0.5% (w/v) Bovine Serum Albumin (BSA), 0.2% Triton-X in PBS (washing solution) was added and left on ice overnight to dissolve Matrigel. Organoids were then transferred into multiple CellCarrier-96 Ultra Microplates (PerkinElmer, 6055300) for staining. Blocking was performed in washing solution with 5% donkey serum (Stratech, 017-000-121-JIR), 0.5% (w/v) Bovine Serum Albumin (BSA) (blocking solution) at 4°C overnight. For primary antibody staining, the following antibodies in blocking solution were used at 4 °C for 24-48 hrs: SOX2 (1: 500, Bio-techne, AF2018), SOX9 (1: 500, Sigma, AB5535), E-cadherin (1: 1500, Thermo Fisher Scientific, 13-1900), GFP (1: 500, AbCam, ab13970). After washing off the primary antibodies, the following secondary antibodies in washing buffer were used at 4°C overnight: donkey anti-chick Alexa 488 (1: 2000, Jackson Immune, 703-545-155), donkey anti-rabbit Alexa 488 (1: 2000, Thermo Fisher Scientific, A21206), donkey anti-rabbit Alexa 594 (1: 2000, Thermo Fisher Scientific, A-21207), donkey anti-goat Alexa 594 (1: 2000, Thermo Fisher Scientific, A-11058), donkey anti-rat Alexa 647 (1: 2000, Jackson Immune, 712-605-153). The following day, DAPI (Sigma, D9542) staining was performed in washing solution at 4°C for 30 min. After two washes with PBS, 97% (v/v) 2’-2’-thio-diethanol (TDE, Sigma, 166782) in PBS was used for mounting. Confocal z stacks were acquired using Leica SP8 at an optical resolution of 1024 × 1024 using a 40x lens. Images were processed using ImageJ (version 2.0.0).

### Immunofluorescence for human fetal lung sections

Human fetal lung sections were rinsed PBS twice and were permeabilized with 0.3% Triton-X in PBS for 10 min. After three PBS washes were carried out, 5% donkey serum (Stratech, 017-000-121-JIR), 0.5% (w/v) Bovine Serum Albumin (BSA), 0.1% Triton-X in PBS (blocking solution) was used for blocking at RT for 1 hr, or at 4°C overnight. Primary antibodies, including SOX2 (1: 500, Bio-techne, AF2018), SOX9 (1: 500, Sigma, AB5535), E-cadherin (1: 1500, Thermo Fisher Scientific, 13-1900) and CD44 (1:100, Thermo Fisher Scientific, 14-0441-82), in blocking solution were used at 4°C overnight. After washing off the primaries, the secondary antibodies in 0.1% Triton-X in PBS were used at 4°C overnight. The following day, DAPI (Sigma, D9542) staining was performed in 0.1% Triton-X in PBS at RT for 20 min. After three washes in PBS, Fluoromount− Aqueous Mounting Medium (Sigma, F4680) was used for mounting. Confocal z stacks or single planes were acquired using Leica SP8 at an optical resolution of 1024 × 1024 at 40x. Single plane images are shown. Images were processed using ImageJ (version 2.0.0).

### Lentiviral production

HEK293T cells were grown in 10-cm dishes to 80% confluency before transfection with the lentiviral vector (10 µg) with packaging vectors including pMD2.G (3 µg, Addgene plasmid # 12259), psPAX2 (6 µg, Addgene plasmid # 12260) and pAdVAntage (3 µg, E1711, Promega) using Lipofectamine 2000 Transfection Reagent (11668019, Thermo Fisher Scientific) according to manufacturer’s protocol. After 16 hrs, medium was refreshed. Supernatant containing lentivirus was harvested at 24 hrs and 48 hrs after medium refreshing and pooled together. Supernatant was centrifuged at 300g to remove cell fragments and passed through 0.45 µm filter. Lentivirus containing >10 kb length insert (inducible CRISPRi) was concentrated using AVANTI J-30I centrifuge (Beckman Coulter) with JS-24.38 swing rotor at 72000g for two hours at 4°C and pellets were dissolved in 200 µl PBS. Other Lentivirus, including individual gRNAs, constructively active β-catenin and SOX9 inducible overexpression, were concentrated using Lenti-X− Concentrator (631232, Takara) and pellets were dissolved in 400 µl PBS.

### gRNA library cloning

gRNA plasmid (Addgene #167936) was linearized with BbsI-HF restriction enzyme overnight at 37 °C. Linearized plasmid was gel purified with QIAquick® Gel Extraction Kit (28704, Qiagen).

The top 5 gRNA sequences for each targeted gene and #0-#32 non-targeting control gRNA sequences were selected from (Gilbert et al, 2016). The gRNA library sequences are listed in **Supplementary Table 1**. 25 nucleotide homology sequences flanking the BbsI cleavage sites of the gRNA plasmid were added at 5’ and 3’ ends of the gRNA sequences for cloning. Oligos were purchased as oPools− Oligo Pools from IDT at 10 pmol per oligo scale.

1.0 pmol of oligo pool and 0.005 pmol linearized plasmid were mixed with NEBuilder® HiFi DNA Assembly Master Mix (E2621, New England BioLabs) according to the manufacturer’s instructions. The mixture was incubated at 50°C for 1 hr and cooled down on ice. Three transformation reactions were performed with 2 µl of the reaction mixture in 50 µl Stellar− Competent Cells (636763, Takara) each according to the manufacturer’s manual. A small fraction of the competent cells was plated to calculate the amount needed for 500X representation per gRNA as seeding in Maxiprep. 10 colonies were randomly picked, minipreped (27104, Qiagen) and sequenced. No repetitive gRNA sequence or backbone sequence were observed. At the same time, same amount of linearized vector was transformed in the same way for background control (colony number < 5% of the library construction transformation). gRNA library pool plasmid was harvested using EndoFree Plasmid Maxi Kit (12362, Qiagen). gRNA library quality was subsequently checked by next generation sequencing (NGS). Primer sequences used are listed in **Supplementary Table 8**.

### Lentiviral production for CRISPRi gRNA library

The gRNA library packaging was slightly modified from a previous method reported (Tian et al., 2019). Two 15-cm dishes with 80-90% confluent HEK293T cells were used for transfection. 10 µg CRISPRi gRNA library plasmid, pMD2.G (2.2 µg, Addgene plasmid # 12259), psPAX2 (4 µg, Addgene plasmid # 12260) and pAdVAntage (1.6 µg, E1711, Promega) were diluted into 2mL Opti-MEM I Reduced Serum Medium (51985034, Thermo Fisher Scientific). 250 µl Lipofectamine 2000 Transfection Reagent (11668027, Thermo Fisher Scientific) was added into 2mL Opti-MEM and incubated at room temperature for 5 min. The diluted DNA mixture was added to the diluted Lipofectamine solution, inverted several times to mix, and incubated at room temperature for 15 min. The transfection mixture was then distributed dropwise to each 15-cm dish with HEK293T cells and mixed well. The next morning, medium was refreshed with DMEM/F12 with 10% FBS. Two days later, the medium was harvested, spun down and filtered through 0.45 µm filters. Lenti-X− Concentrator (631231, Takara) was added to the supernatant at 1: 3 (v/v) ratio. After 48 hrs, lentivirus was pelleted at 1500g for 45 min at 4 °C. Lentivirus pellets were re-suspended with 20 ml of self-renewing medium with ROCK inhibitor (ROCKi, Y-27632, 10 µM, 688000, Merck), aliquoted and stored in -80 °C before use.

### CRISPRi screen

Lentivirus titration was first performed to determine the amount of CRISPRi gRNA lentivirus needed for the screen. BRC1915 and HDBR-13393 inducible CRISPRi parental organoid lines were dissociated with TrypLE (Thermo Fisher Scientific, 12605028) at 37°C for 10-20 min (with pipette trituration every 5 min). Organoid single cells were counted and re-suspended to achieve 100K cells/500 µl of self-renewing medium with ROCKi in each 24-well plate well. A serial gradient of the CRISPRi gRNA library lentivirus was added and incubated at 37 °C overnight. The next morning, organoid cells were collected, washed twice with PBS and seeded back to 2x 24 well plate well/condition. Flow cytometry analysis was performed after 3 days to determine the amount of virus needed to reach 10-30% of infection rate per 100K cells per 500 µl of medium.

BRC1915 and HDBR-13393 inducible CRISPRi parental organoid lines were expanded and dissociated with TrypLE at 37°C for 10-20 min (with pipette trituration every 5 min). 4.2 million and 5 million organoid cells from each parental organoid lines were used for infection respectively. Organoid cells were counted and re-suspended at 100K cells per 500 µl of medium. The infection was performed in a 15-cm dish with the appropriate amount of lentivirus determined by titration. Organoid cells were collected the next morning, washed and seeded back to Matrigel with 50K cells/50 µl Matrigel dome in 6 well plates with 8 Matrigel domes per well. Self-renewing medium with ROCKi was supplemented for 3 days before organoids were dissociated with TrypLE and prepared for cell sorting. The TagRFP^+^EGFP^+^ double positive population was harvested. Organoid cells from BRC1915 line were divided into 3 parts, each with 983x cell representation per gRNA. One part was snap frozen for gRNA abundance analysis and the two remaining parts were separated as technical replicates and seeded back into Matrigel with 10 K cells/50 µl Matrigel dome in 6 well plates with 8 Matrigel domes per well. Organoid cells from HDBR-13393 were divided into 2 parts, each with 480x representation per gRNA. One part was snap frozen for gRNA abundance analysis and the other part was seeded back to Matrigel with 10 K cells/50 µl Matrigel dome in 6 well plates with 8 Matrigel domes per well. Organoid cells were cultured in self-renewing medium with ROCKi for 8 days before Dox (Doxycycline hyclate, 2 µg/ml, Merck, D9891) and TMP (Trimethoprim, 10 µM, Merck, 92131) were administered and cultured for 2 weeks further.

Organoids were passaged (physically split by breaking into pieces) twice during this time. Organoid cells were then dissociated into single cells and prepared for flow cytometry cell sorting. TagRFP^high^EGFP^+^ and TagRFP^low^EGFP^+^ cells were sorted, harvested separately and snap frozen for gRNA abundance analysis.

Genomic DNA was isolated with Macherey-Nagel− NucleoSpin− Blood kit (740951.50, Macherey-Nagel) according to manufacturer’s instruction. 2 µg of genomic DNA for BRC1915 organoid line and 1 µg genomic DNA for HDBR-N 13393 organoid line was used for PCR amplification of gRNA for NGS. NEBNext Ultra II Q5 Master Mix (M0544S, New England BioLabs) and primers (**Supplementary Table 8**) were used for amplification with 29 cycles. PCR products were gel purified with Monarch® DNA Gel Extraction Kit (T1020S, New England BioLabs) according to manufacturer’s instructions and sent for NGS at the Gurdon Institute NGS core using an Illumina HiSeq 1500.

### CRISPRi screen analysis

Guide abundances were quantified by mapping reads present in sequencing data to the CRISPR library sequences. The initial nucleotide was omitted from both library sequences and reads, as the initial base of a read is often not certain. Mapped read counts are available in **Supplementary Table 2**. MAGeCK (Li et al., 2014) was used to determine significantly enriched or depleted genes. As MAGeCK expects the majority of genes targeted to have no effect, the 33 non-targeting guides were bootstrapped to form 102 synthetic genes by drawing 5 unique random guides from the non-targeting group. MAGeCK version 0.5.9.3 was run with “--remove-zero both” and “--remove-zero-threshold 1” options.

### CRISPRi screen hit validation

The 2 most effective gRNA sequences in the screen for *MYBL2, IRF6, ARID5B* and *ZBTB7B* were selected (**Supplementary Table 8**). These gRNAs were individually cloned into the gRNA plasmids and packaged into lentivirus as described above.

3 independent inducible CRIPSRi parental organoid lines were transduced individually with specific gRNAs. TagRFP and EGFP double positive organoid cells were sorted and seeded in Matrigel with 2000-3000 cells per 50 µl Matrigel (for gRNAs targeting the same gene, the same number of cells were used from each parental organoid line). Organoid cells were cultured with self-renewing medium plus ROCKi for 8 days before Dox and TMP were administered for 5 days. One third of the organoids was harvested for mRNA isolation to check for targeted gene knockdown. RNA extraction was performed using RNeasy Mini Kit (74104, Qiagen) with RNase-Free DNase Set (Qiagen) according to the manufacturer’s instructions. Reverse transcription was performed using MultiScribe− Reverse Transcriptase system (4311235, Thermo Fisher Scientific) according to the manufacturer’s instructions. qRT-PCR primers for each target genes are listed in **Supplementary Table 8**. One third of the organoids was used for EdU incorporation assay with 1-hour pulse labelling, fixed and staining with Click-iT− EdU Cell Proliferation Kit for Imaging (C10340, Thermo Fisher Scientific) according to the manufacturer’s instructions. The rest of the organoids were passaged at 1 to 3 ratio till Passage #5, or the line was lost.

### *SOX2* and *SOX9* knockdown RNA-Seq

BRC1915 and HDBR-N 13393 inducible CRISPRi parental organoid lines were used for these experiments. For each parental line, NT gRNAs (#1, #2), *SOX2* gRNAs (#2, #8), and *SOX9* gRNAs (#1, #7) were introduced via lentivirus transduction (gRNA sequences in **Supplementary Table 8**). TagRFP and EGFP double positive cells were sorted, re-plated and expanded. Organoids were treated with Dox and TMP for 5 days, before harvesting for RNA extraction, which was performed using RNeasy Mini Kit (Qiagen) with RNase-Free DNase Set (Qiagen) according to the manufacturer’s instructions.

RNA was quantified by NanoDrop (Thermo Fisher Scientific) and Tape Station (Agilent) with High Sensitivity RNA Screen Tape (5067-5579, Agilent). Half of the RNA was used for cDNA synthesis and qRT-PCR analysis to validate *SOX2* and *SOX9* knockdown effect. The other half of the RNA was sent for Eukaryotic RNA-seq library preparation and sequencing (PE150) at Novogene (Cambridge, UK).

### RNA-Seq data analysis

Raw sequencing data was aligned to human genome (hg38) using STAR aligner (v 2.7.1a). Sequencing reads were counted using FeatureCount (v 2.0.0). DESeq2 was used for differential gene expression analysis. Differentially expressed (DE) gene lists were generated using DESeq2 packaged with a cut-off of fold change ≥ 2 and FDR ≤ 0.05. *SOX2* KD, *SOX9* KD or NT control with different gRNAs of the same organoid line were first grouped for DE gene analysis. DE genes shared by two independent organoid lines were taken as the most robust DE gene list for downstream analysis. Differentially expressed genes were used for GO analysis with ShinyGO v0.61 (http://bioinformatics.sdstate.edu/go/).

### Comparing SOX9 targets signature with human fetal lung scRNA-Seq data

Up-regulated and down-regulated genes in SOX9 knockdown DE gene list and SOX9 direct transcriptional target list were used to score epithelial cells from 9-11 pcw human fetal lungs (He et al., 2022), using Scanpy’s tl.score_genes function.

### SOX9, ETV4 and ETV5 TaDa

Four organoid lines (HDBR-N 13393, BRC1915, BRC1929, BRC1938) were used for SOX9, ETV4 and ETV5 TaDa experiments. For each organoid line, 4 ×10^5^ organoid single cells were prepared as aforementioned method and re-suspended in 500 µl self-renewing medium with ROCKi in one 24-well plate well. 40 µl of SOX9, ETV4, ETV5 or Dam-only TaDa lentivirus was added per well and incubated overnight. The next morning, organoid single cells were harvested and washed twice with PBS before being re-suspended with 200 µl of Matrigel and seeded back to four 24-well plate wells. After 48-72 hrs, organoids were harvested for genomic DNA isolation.

### DamID-seq

DamID samples were processed as described previously and prepared for Illumina sequencing with an adapted TruSeq protocol (Marshall et al., 2016). All sequencing runs were performed as single end 50 bp reads using an Illumina HiSeq 1500, or single end 100 bp reads using an Illumina NovaSeq 6000 from the Gurdon Institute NGS Core facility.

### DamID-seq data processing

Analysis of the raw fastq files was performed with the damidseq pipeline script (Marshall and Brand, 2015) that maps reads to an indexed bowtie2 genome, bins into GATC-fragments according to GATC-sites and normalises reads against a Dam-only control. For our data, the fastq files were mapped to the GRCh38 genome assembly (hg38). The different patient-derived organoid samples were treated as individual replicates. For each replicate the signal from the Dam-fusion samples was normalized against its own Dam-only sample with a modified version of the damidseq_pipeline (RPM normalization, 300 bp bin size). Binding intensities were quantile normalised across all replicates for the same Dam-fusion and subsequently averaged and inversed (“unlog”). Files were converted to the bigwig file format with bedGraphToBigWig (v4) for visualization with the Integrative Genomics Viewer (IGV) (v2.4.19).

### Peak calling and analysis

Macs2 (v2.1.2) was used to call broad peaks for every Dam-fusion and Dam-only pair using the bam files generated by the damidseq_pipeline. Peaks were merged when they were within 1kb from each other with the merge-function from bedtools (v2.26.0). Peaks were selected when present in all replicates for a particular experimental condition with a FDR<10^−50^. Overlapping genomic features were identified by uploading the peak coordinates to the GREAT online platform (http://great.stanford.edu/public/html/). Motif analysis was performed using HOMER (v 4.11) analysis with findMotifsGenome.pl function. Genomic occupancy feature annotation was performed using HOMER (v 4.11) analysis with annotatePeaks.pl function. Correlation analysis of different TaDa results were performed using deepTools (v 3.5.0) multiBigwigSummary function.

### In situ hybridization chain reaction (in situ HCR)

*In situ* HCR v3.0 was performed according to the manufacturer’s procedure (Molecular Instruments). A set of DNA HCR probes were designed according to the protocol (Choi et al., 2018) and the HCR amplifiers with buffers were purchased from Molecular Instruments. Sequence information for the DNA HCR probes for *SOX9, ETV4, ETV5, LGR5* and *NOTUM* mRNA targets are listed in **Supplementary Table 9**. In brief, 14 μm frozen human tissues sections were rinsed in nuclease-free water and pre-hybridized for 10 min in 37 °C humidified chamber. Each DNA HCR probe was diluted to 2 pmol and incubated with the tissues at 37 °C overnight. After a series of washing in probe wash buffer, the tissue slices were incubated with 6 pmol of HCR amplifiers at room temperature overnight for amplification reaction. The HCR amplifier comprised of two hairpins, B1 and B2, labeled with Alexa 546 and 647, respectively, were snap-cooled separately at 3 µM before adding to the tissue sections. After removing excess hairpins in 5X SSCT (sodium chloride sodium citrate) buffer, nuclei were stained with DAPI. Images were collected on an Leica SP8 confocal microscopy.

### Western Blot

Organoids were harvested and washed twice with Advanced DMEM/F12 and twice with PBS, before organoids were re-suspended in 100 µl-200 µl of RIPA buffer with protease inhibitor (1x, Thermo Fisher Scientific, 78440) added. Organoid suspension was incubated for 30 minutes on ice, with strong vortex every 5 minutes. Cell pieces and debris were removed by centrifugation at 13000 rpm. Supernatant was harvested. Protein concentration was measured by BCA assay (Thermo Fisher Scientific). Equal amount of each protein sample was mixed with Sample Buffer (Bio-rad) and beta-mercaptoethanol (Merck, M6250) according to manufacturer’s protocol. Mixture was heated at 95°C for 5 minutes and cooled down to room temperature.

Samples were separated on a 4%–12% SDS-PAGE and transferred to nitrocellulose membranes. Proteins were detected by incubation with primary antibodies, SOX9 (1: 1000, Merck, AB5535) and β-Actin (1: 5000, Merck, A3854) at 4 °C overnight and subsequently secondary antibodies, donkey anti-Mouse IRDye 800CW (1: 1000, AbCam, ab216774) and donkey anti-Rabbit IRDye 680RD (1: 1000, AbCam, ab216779). Protein bands were visualised using Li-Cor Odyssey system.

### Molecular cloning

Inducible CRIPSRi and gRNA plasmids (Addgene: #167935 and #167936 respectively) were previously deposited at Addgene. The TaDa Dam-only plasmid was generated by In-fusion cloning of mNeonGreen and *E*.*coli* Dam methylase into a linearized plasmid pHR-SFFV-dCas9-BFP-KRAB (a gift from Stanley Qi & Jonathan Weissman, Addgene plasmid # 46911), with MluI and SbfI restriction enzymes. SOX9, ETV4 and ETV5 TaDa plasmids (N-terminal Dam fusion) were generated by Infusion cloning of the target gene sequence (amplified from cDNA library) into the linearized TaDa Dam-only plasmid with NotI restriction enzyme. Constitutively The constitutively activated β-catenin overexpression plasmid was generated by Infusion cloning of deltaN90 β-catenin (amplified from cDNA library) and mNeonGreen into the linearized inducible CRISPRi plasmid with XhoI and BamHI restriction enzymes.

---

**Figure S1.**
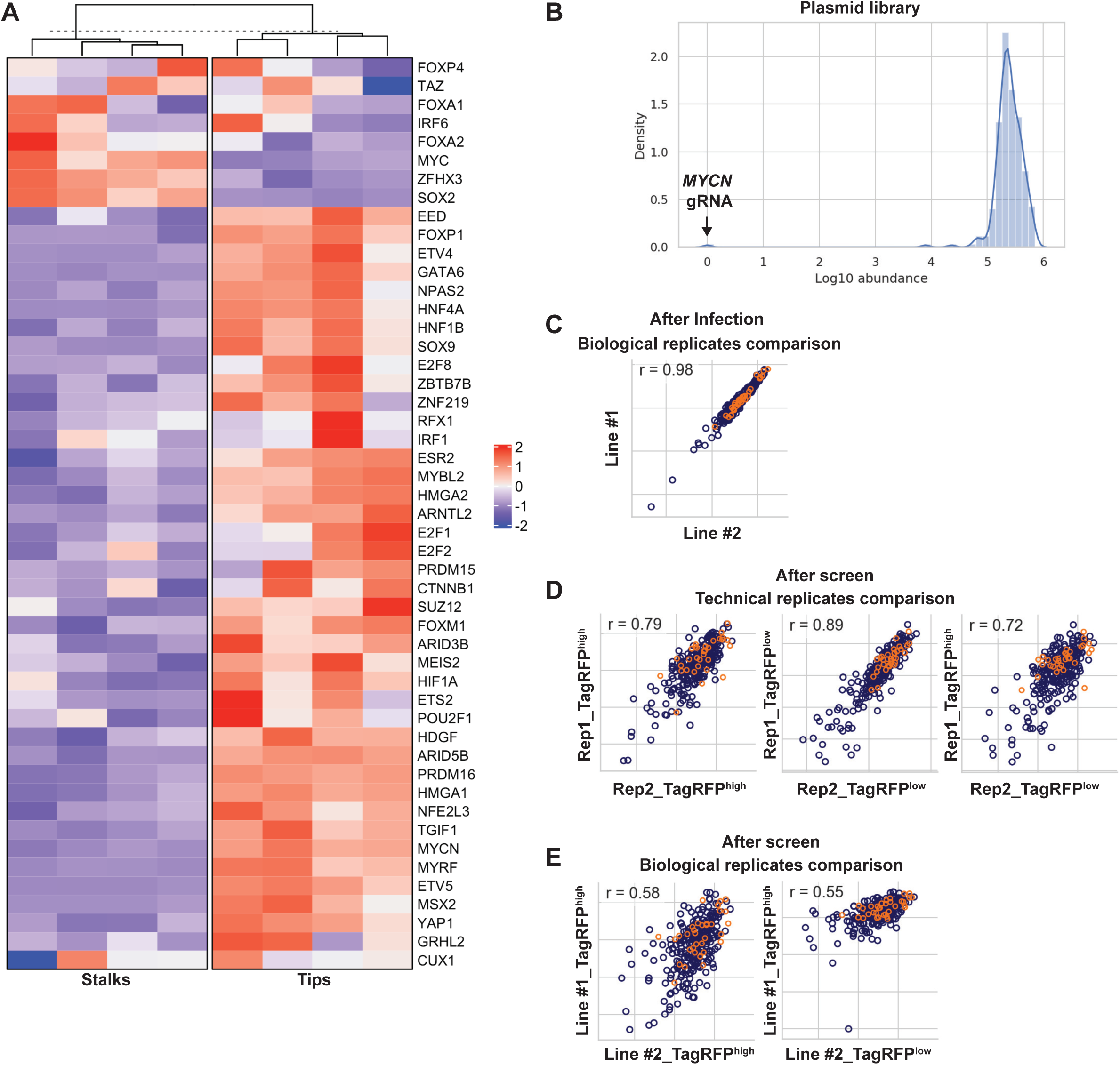
CRISPRi screen quality control. (**A**) Expression levels heatmap of the selected transcription factors in the developing human fetal lung tip progenitor and stalk cells. Data from Nikolic et al., 2017. (**B**) gRNA abundance distribution of the CRISPRi library after cloning into the plasmid vector. One gRNA targeting *MYCN* was missing; likely due to a gRNA synthesis issue. (**C**) Pearson correlation of gRNA abundance between different samples indicated in axes. Between two independent CRISPRi parental lines 3 days after lentiviral transduction. R = 0.98 indicated great consistency of lentiviral transduction. (**D**) Pearson correlation of gRNA abundance between technical replicates (Rep1 and Rep2). Great consistency was observed between TagRFP^high^ and TagRFP^low^ populations. (**E**) Pearson correlation of gRNA abundance between biological replicates. A lower correlation was observed reflecting the variation of human tissue samples. Orange circles in (**C-E**) representing non-targeting control gRNAs.

**Figure S2.**
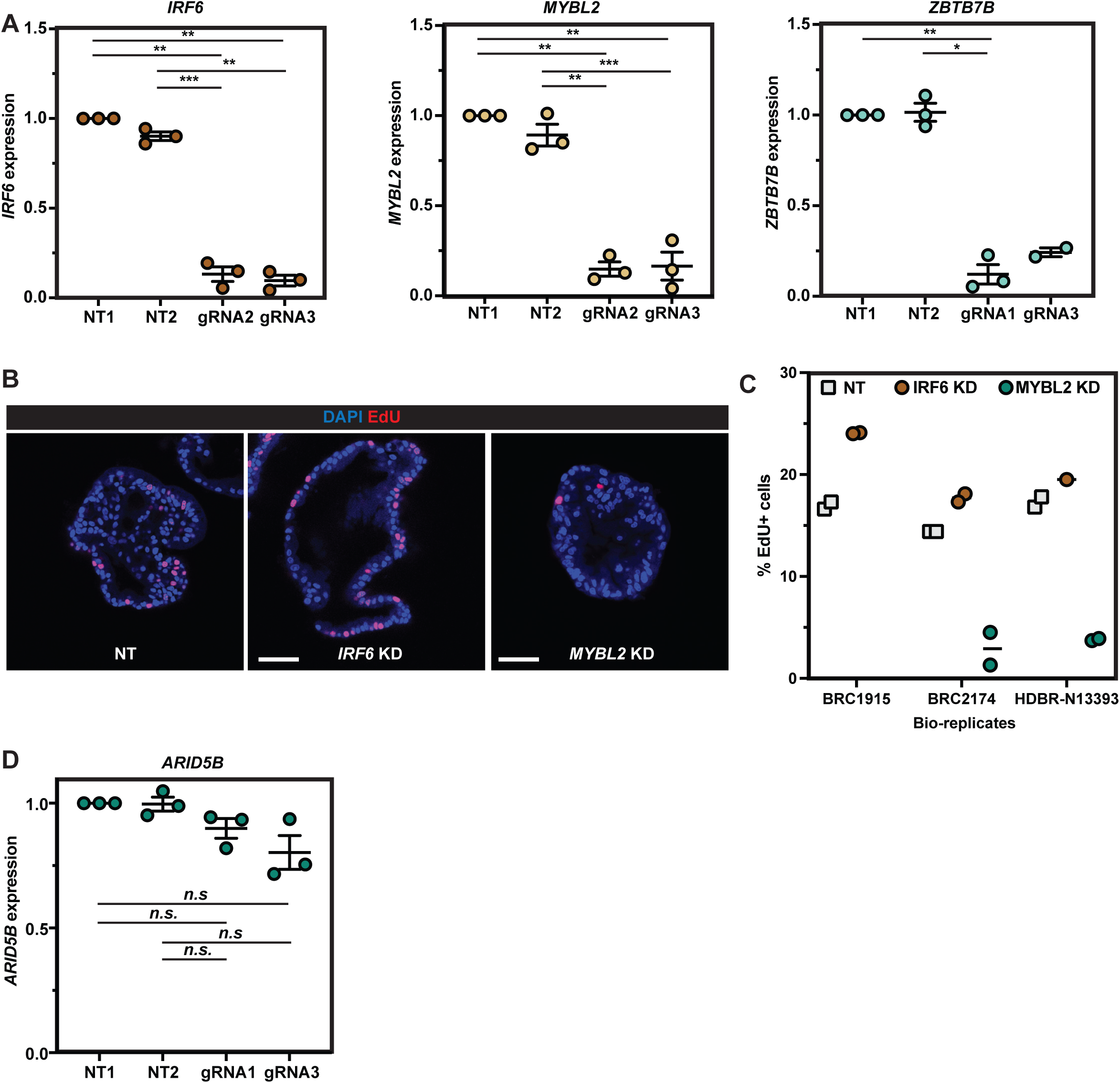
Validation of the CRISPRi screen results. (**A**) qRT-PCR results showing the targeted genes (*IRF6, MYBL2, ZBTB7B*) were efficiently knocked down by the inducible CRISPRi system using the gRNAs selected from the CRISPRi gRNA library. (**B**) Representative EdU staining images of non-targeting gRNA control and *IRF6* or *MYBL2* knockdown experiments. (**C**) Quantification of the percentage of EdU^+^ cells in each of two parental organoid lines used with non-targeting control, *IRF6* knockdown and *MYBL2* knockdown. n = 1649, 1705, 3548 cells were scored for NT controls. n = 2517, 950, and 1313 cells were scored for *IRF6* gRNAs. n = 1098 and 1306 cells were scored for *MYBL2* gRNAs. (**D**) qRT-PCR results showing *ARID5B* was not knocked down by the inducible CRISPRi system using the gRNAs selected from the CRISPRi gRNA library. Error bars: mean ± SEM. Statistical analysis was using the two-tailed paired T-test. p values are reported as follows: *p < 0.05, **p < 0.01, ***p < 0.001 and n.s. non-significant. N=3 organoid lines (biological replicates) used for each panel.

**Figure S3.**
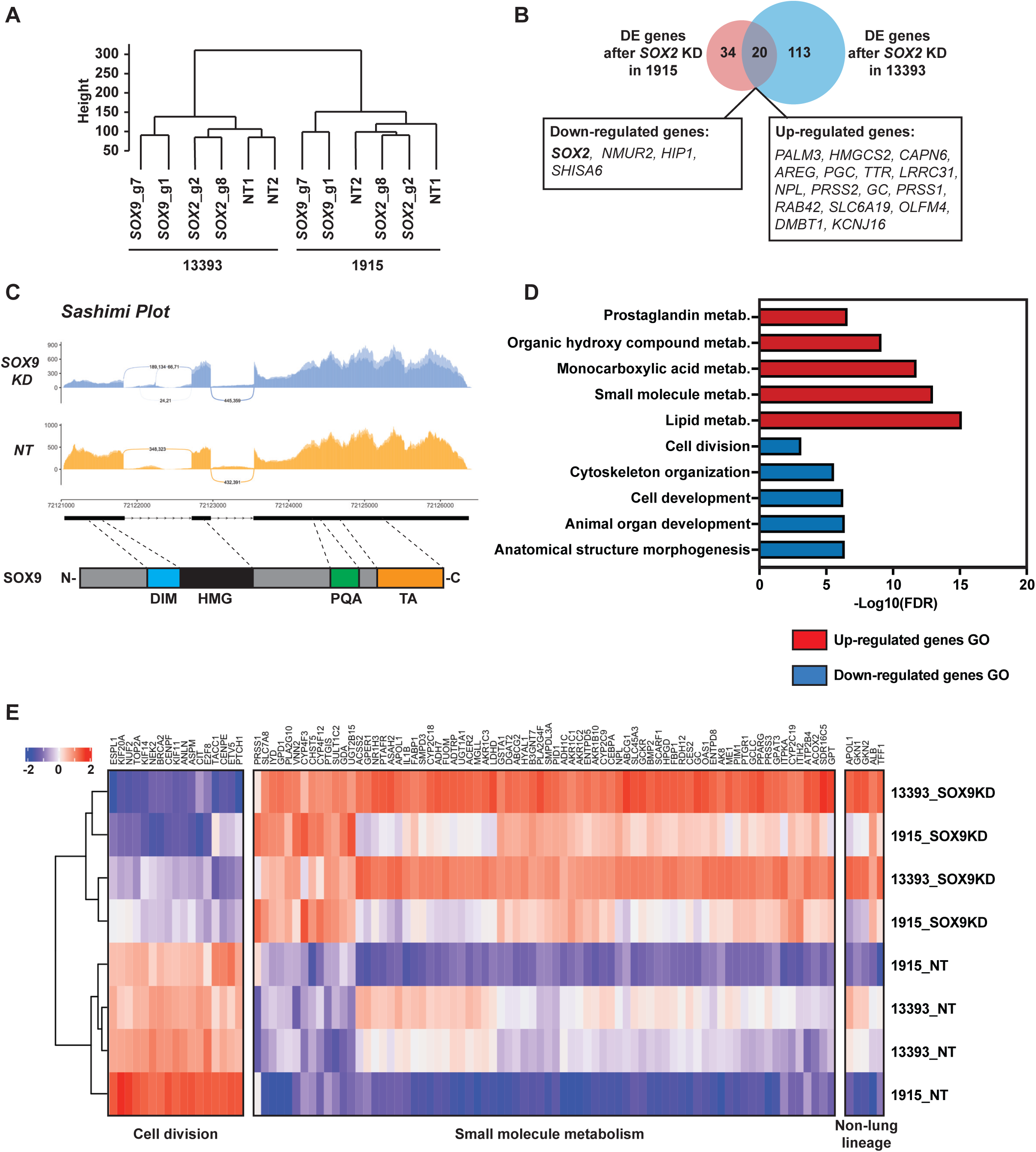
*SOX2* and *SOX9* knockdown resulted in different transcriptome changes. (**A**) Unsupervised hierarchical clustering of non-targeting control, *SOX2* knockdown and *SOX9* knockdown RNA-Seq results. (**B**) Venn diagram showing minimal overlap of differentially expressed genes after *SOX2* knockdown in two different parental organoid lines. Overlapping DE genes were labelled in boxes. (**C**) Sashimi plot to visualise splicing junction of NT control and SOX9 KD. Upper panel: Sashimi plot was used to visualise splicing junction information in non-targeting gRNA control and *SOX9* knockdown groups. Junctional reads between intron #1 and exon #2 were only observed in *SOX9* knockdown groups and not in non-targeting gRNA control groups. Lower panel: major SOX9 domains in relation to the *SOX9* genomic locus. Exon #1 contains DIM and part of the HMG domain. DIM, dimerization domain. HMG, high-mobility group domain. PQA, proline-glutamine-alanine repeats domain. TA, transactivation domain. (**D**) Selected GO enrichment in DE genes after SOX9 knockdown. (**E**) Heatmap of gene expression from representative GO terms: cell division and small molecule metabolism together with gene expression of upregulated non-lung lineage genes.

**Figure S4.**
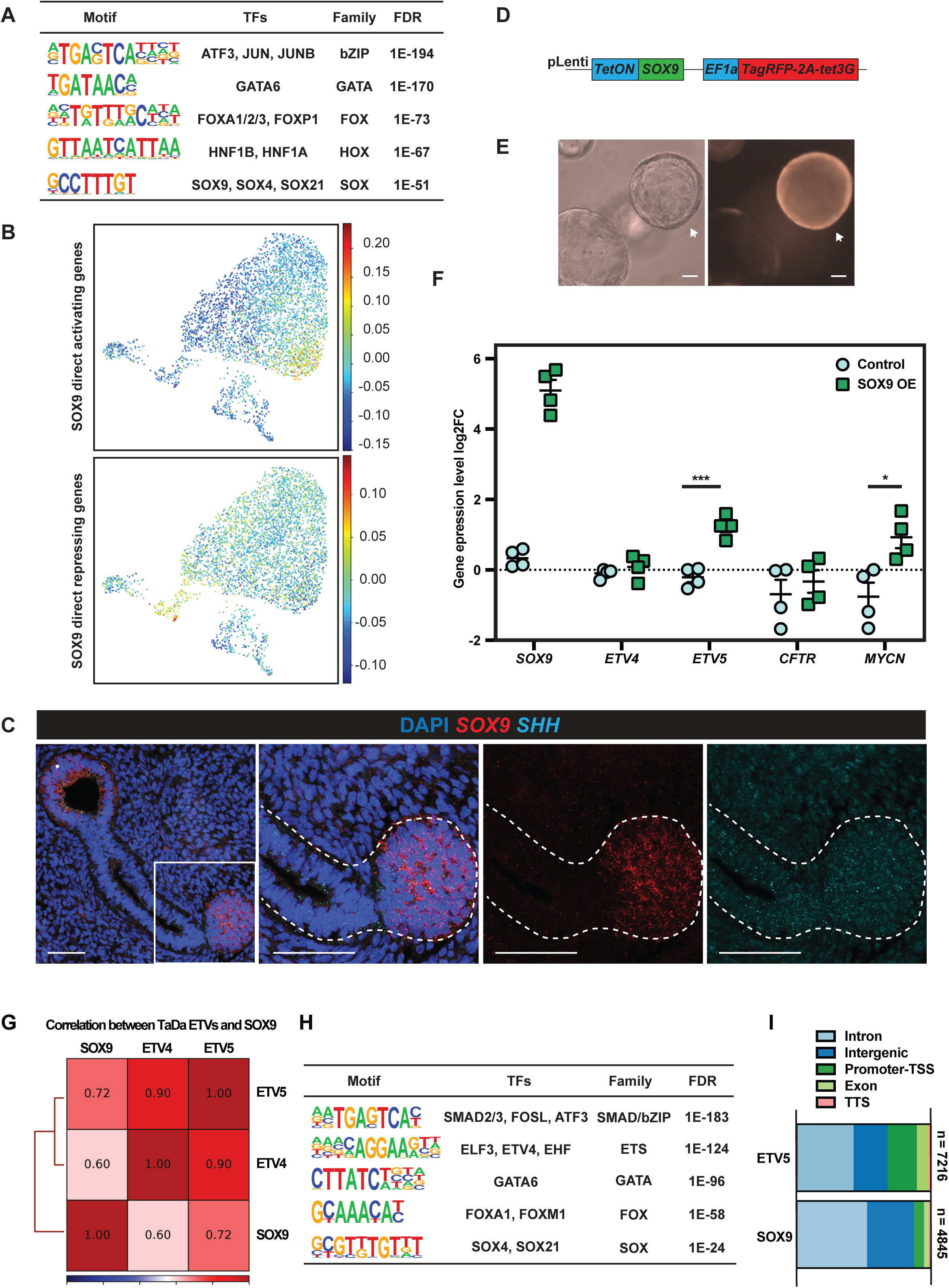
SOX9 directly activates tip cell genes and represses secretory cell genes. (**A**) Summary of enriched TF binding motifs in SOX9 TaDa peaks. The SOX motif was enriched, indicating the SOX9 TaDa faithfully identified SOX9 binding sites across the genome. (**B**) SOX9 direct transcriptional targets enrichment in human fetal lung scRNA-Seq data. All of SOX9 direct transcriptional targets were used for scoring. Similar to Fig. 2H, SOX9 directly-activated targets were enriched in tip progenitor cells (upper panel) whereas SOX9 directly-repressed targets were enriched in secretory cell lineages (lower panel). (**C**) *SHH* was co-expressed with *SOX9* in human fetal lung tip progenitor cells. S*OX9* in red and *SHH* in cyan. **(D)** Lentiviral construct design for overexpressing SOX9 in human fetal lung progenitor cells. **(E)** Representative images showing organoid morphology does not change after 3 days of SOX9 overexpression. SOX9 overexpressed organoid indicated with arrow. (**F**) qRT-PCR results showing that SOX9 overexpression led to *ETV5* and *MYCN* transcription being significantly upregulated, however, *ETV4* and *CFTR* were not changed. Two-tailed Student’s T-tests were performed. p values are reported as follows: *p < 0.05; ***p < 0.001. (**G**) Pearson correlation of SOX9, ETV4 and ETV5 TaDa. ETV4 and ETV5 TaDa exhibited great consistency. (**H**) Motifs enriched in ETV5 TaDa peaks. The ETS binding motif was highly enriched. (**I**) Genomic occupancy annotated features for SOX9 and ETV5 peaks. Scale bars denote 50 µm (C) and 100 µm (E).

## Notes

### Competing Interest Statement

The authors have declared no competing interest.

## REFERENCES

Alanis, D.M., Chang, D.R., Akiyama, H., Krasnow, M.A., Chen, J., 2014. Two nested developmental waves demarcate a compartment boundary in the mouse lung. Nat Commun 5, 3923. doi:10.1038/ncomms4923

Botti, E., Spallone, G., Moretti, F., Marinari, B., Pinetti, V., Galanti, S., De Meo, P.D., De Nicola, F., Ganci, F., Castrignanò, T., Pesole, G., Chimenti, S., Guerrini, L., Fanciulli, M., Blandino, G., Karin, M., Costanzo, A., 2011. Developmental factor IRF6 exhibits tumor suppressor activity in squamous cell carcinomas. Proc. Natl. Acad. Sci. U.S.A. 108, 13710–13715. doi:10.1073/pnas.1110931108

Bowden, A.R., Morales-Juarez, D.A., Sczaniecka-Clift, M., Agudo, M.M., Lukashchuk, N., Thomas, J.C., Jackson, S.P., 2020. Parallel CRISPR-Cas9 screens clarify impacts of p53 on screen performance. Elife 9, 2430. doi:10.7554/eLife.55325

Chang, D.R., Martinez Alanis, D., Miller, R.K., Ji, H., Akiyama, H., McCrea, P.D., Chen, J., 2013. Lung epithelial branching program antagonizes alveolar differentiation. Proc. Natl. Acad. Sci. U.S.A. 110, 18042–18051. doi:10.1073/pnas.1311760110

Cheetham, S.W., Gruhn, W.H., van den Ameele, J., Krautz, R., Southall, T.D., Kobayashi, T., Surani, M.A., Brand, A.H., 2018. Targeted DamID reveals differential binding of mammalian pluripotency factors. Development 145, dev170209. doi:10.1242/dev.170209

Chen, Y.-W., Huang, S.X., de Carvalho, A.L.R.T., Ho, S.-H., Islam, M.N., Volpi, S., Notarangelo, L.D., Ciancanelli, M., Casanova, J.-L., Bhattacharya, J., Liang, A.F., Palermo, L.M., Porotto, M., Moscona, A., Snoeck, H.-W., 2017. A three-dimensional model of human lung development and disease from pluripotent stem cells. Nat. Cell Biol. 19, 542–549. doi:10.1038/ncb3510

Cho, S.W., Kim, S., Kim, J.M., Kim, J.-S., 2013. Targeted genome engineering in human cells with the Cas9 RNA-guided endonuclease. Nat. Biotechnol. 31, 230–232. doi:10.1038/nbt.2507

Cong, L., Ran, F.A., Cox, D., Lin, S., Barretto, R., Habib, N., Hsu, P.D., Wu, X., Jiang, W., Marraffini, L.A., Zhang, F., 2013. Multiplex genome engineering using CRISPR/Cas systems. Science 339, 819–823. doi:10.1126/science.1231143

Danopoulos, S., Alonso, I., Thornton, M.E., Grubbs, B.H., Bellusci, S., Warburton, D., Alam Al, D., 2018. Human lung branching morphogenesis is orchestrated by the spatiotemporal distribution of ACTA2, SOX2, and SOX9. Am. J. Physiol. Lung Cell Mol. Physiol. 314, L144–L149. doi:10.1152/ajplung.00379.2017

Doench, J.G., Fusi, N., Sullender, M., Hegde, M., Vaimberg, E.W., Donovan, K.F., Smith, I., Tothova, Z., Wilen, C., Orchard, R., Virgin, H.W., Listgarten, J., Root, D.E., 2016. Optimized sgRNA design to maximize activity and minimize off-target effects of CRISPR-Cas9. Nat. Biotechnol. 34, 184–191. doi:10.1038/nbt.3437

Fulco, C.P., Munschauer, M., Anyoha, R., Munson, G., Grossman, S.R., Perez, E.M., Kane, M., Cleary, B., Lander, E.S., Engreitz, J.M., 2016. Systematic mapping of functional enhancer–promoter connections with CRISPR interference. Science. doi:10.1126/science.aag2445

Fulco, C.P., Nasser, J., Jones, T.R., Munson, G., Bergman, D.T., Subramanian, V., Grossman, S.R., Anyoha, R., Doughty, B.R., Patwardhan, T.A., Nguyen, T.H., Kane, M., Perez, E.M., Durand, N.C., Lareau, C.A., Stamenova, E.K., Aiden, E.L., Lander, E.S., Engreitz, J.M., 2019. Activity-by-contact model of enhancer–promoter regulation from thousands of CRISPR perturbations. Nat Genet 51, 1664–1669. doi:10.1038/s41588-019-0538-0

Gasperini, M., Hill, A.J., McFaline-Figueroa, J.L., Martin, B., Kim, S., Zhang, M.D., Jackson, D., Leith, A., Schreiber, J., Noble, W.S., Trapnell, C., Ahituv, N., Shendure, J., 2019. A Genome-wide Framework for Mapping Gene Regulation via Cellular Genetic Screens. Cell 176, 377–390.e19. doi:10.1016/j.cell.2018.11.029

Gerner-Mauro, K.N., Akiyama, H., Chen, J., 2020. Redundant and additive functions of the four Lef/Tcf transcription factors in lung epithelial progenitors. Proc. Natl. Acad. Sci. U.S.A. 117, 12182–12191. doi:10.1073/pnas.2002082117

Gilbert, L.A., Horlbeck, M.A., Adamson, B., Villalta, J.E., Chen, Y., Whitehead, E.H., Guimaraes, C., Panning, B., Ploegh, H.L., Bassik, M.C., Qi, L.S., Kampmann, M., Weissman, J.S., 2014. Genome-Scale CRISPR-Mediated Control of Gene Repression and Activation. Cell 159, 647–661. doi:10.1016/j.cell.2014.09.029

Gilbert, L.A., Larson, M.H., Morsut, L., Liu, Z., Brar, G.A., Torres, S.E., Stern-Ginossar, N., Brandman, O., Whitehead, E.H., Doudna, J.A., Lim, W.A., Weissman, J.S., Qi, L.S., 2013. CRISPR-mediated modular RNA-guided regulation of transcription in eukaryotes. Cell 154, 442–451. doi:10.1016/j.cell.2013.06.044

Guo, W., Keckesova, Z., Donaher, J.L., Shibue, T., Tischler, V., Reinhardt, F., Itzkovitz, S., Noske, A., Zürrer-Härdi, U., Bell, G., Tam, W.L., Mani, S.A., van Oudenaarden, A., Weinberg, R.A., 2012. Slug and Sox9 Cooperatively Determine the Mammary Stem Cell State. Cell 148, 1015–1028. doi:10.1016/j.cell.2012.02.008

He, P., Lim, K., Sun, D., Pett, J.P., Jeng, Q., Polanski, K., Dong, Z., Bolt, L., Richardson, L., Mamanova, L., Dabrowska, M., Wilbrey-Clark, A., Madissoon, E., Tuong, Z.K., Dann, E., Suo, C., Kai’En, I.G., He, X., Barker, R., Teichmann, S.A., Marioni, J.C., Meyer, K.B., Rawlins, E.L., 2022. A human fetal lung cell atlas uncovers proximal-distal gradients of differentiation and key regulators of epithelial fates. bioRxiv 2022.01.11.474933. doi:10.1101/2022.01.11.474933

Hein, R.F.C., Wu, J.H., Tsai, Y.-H., Wu, A., Miller, A.J., Holloway, E.M., Frum, T., Conchola, A.S., Szenker-Ravi, E., Reversade, B., Yan, K.S., Kuo, C.J., Spence, J.R., 2021. R-SPONDIN2+ Mesenchymal Cells Form the Bud Tip Progenitor Niche During Human Lung Development. bioRxiv 2021.04.05.438484. doi:10.1101/2021.04.05.438484

Herriges, J.C., Verheyden, J.M., Zhang, Z., Sui, P., Zhang, Y., Anderson, M.J., Swing, D.A., Zhang, Y., Lewandoski, M., Sun, X., 2015. FGF-Regulated ETV Transcription Factors Control FGF-SHH Feedback Loop in Lung Branching. Dev. Cell 35, 322–332. doi:10.1016/j.devcel.2015.10.006

Horlbeck, M.A., Gilbert, L.A., Villalta, J.E., Adamson, B., Pak, R.A., Chen, Y., Fields, A.P., Park, C.Y., Corn, J.E., Kampmann, M., Weissman, J.S., 2016. Compact and highly active next-generation libraries for CRISPR-mediated gene repression and activation. Elife 5, 914. doi:10.7554/eLife.19760

Jinek, M., Chylinski, K., Fonfara, I., Hauer, M., Doudna, J.A., Charpentier, E., 2012. A Programmable Dual-RNA–Guided DNA Endonuclease in Adaptive Bacterial Immunity. Science 337, 816–821. doi:10.1126/science.1225829

Joung, J., Konermann, S., Gootenberg, J.S., Abudayyeh, O.O., Platt, R.J., Brigham, M.D., Sanjana, N.E., Zhang, F., 2017. Genome-scale CRISPR-Cas9 knockout and transcriptional activation screening. Nature Protocols 12, 828–863. doi:10.1038/nprot.2017.016

Kotton, D.N., Morrisey, E.E., 2014. Lung regeneration: mechanisms, applications and emerging stem cell populations. Nat. Med. 20, 822–832. doi:10.1038/nm.3642

Labaki, W.W., Han, M.K., 2020. Chronic respiratory diseases: a global view. Lancet Respir Med 8, 531–533. doi:10.1016/S2213-2600(20)30157-0

Laughney, A.M., Hu, J., Campbell, N.R., Bakhoum, S.F., Setty, M., Lavallée, V.-P., Xie, Y., Masilionis, I., Carr, A.J., Kottapalli, S., Allaj, V., Mattar, M., Rekhtman, N., Xavier, J.B., Mazutis, L., Poirier, J.T., Rudin, C.M., Pe’er, D., Massagué, J., 2020. Regenerative lineages and immune-mediated pruning in lung cancer metastasis. Nat. Med. 26, 259–269. doi:10.1038/s41591-019-0750-6

Li, L., Feng, J., Zhao, S., Rong, Z., Lin, Y., 2021. SOX9 inactivation affects the proliferation and differentiation of human lung organoids. Stem Cell Res Ther 12, 1–12. doi:10.1186/s13287-021-02422-6

Mali, P., Yang, L., Esvelt, K.M., Aach, J., Guell, M., DiCarlo, J.E., Norville, J.E., Church, G.M., 2013. RNA-Guided Human Genome Engineering via Cas9. Science. doi:10.1126/science.1232033

Mandegar, M.A., Huebsch, N., Frolov, E.B., Shin, E., Truong, A., Olvera, M.P., Chan, A.H., Miyaoka, Y., Holmes, K., Spencer, C.I., Judge, L.M., Gordon, D.E., Eskildsen, T.V., Villalta, J.E., Horlbeck, M.A., Gilbert, L.A., Krogan, N.J., Sheikh, S.P., Weissman, J.S., Qi, L.S., So, P.-L., Conklin, B.R., 2016. CRISPR Interference Efficiently Induces Specific and Reversible Gene Silencing in Human iPSCs. Cell Stem Cell 18, 541–553. doi:10.1016/j.stem.2016.01.022

Marshall, O.J., Brand, A.H., 2015. damidseq_pipeline: an automated pipeline for processing DamID sequencing datasets. Bioinformatics 31, 3371–3373. doi:10.1093/bioinformatics/btv386

Marshall, O.J., Southall, T.D., Cheetham, S.W., Brand, A.H., 2016. Cell-type-specific profiling of protein-DNA interactions without cell isolation using targeted DamID with next-generation sequencing. Nature Protocols 11, 1586–1598. doi:10.1038/nprot.2016.084

Miller, A.J., Hill, D.R., Nagy, M.S., Aoki, Y., Dye, B.R., Chin, A.M., Huang, S., Zhu, F., White, E.S., Lama, V., Spence, J.R., 2018. In Vitro Induction and In Vivo Engraftment of Lung Bud Tip Progenitor Cells Derived from Human Pluripotent Stem Cells. Stem Cell Reports 10, 101–119. doi:10.1016/j.stemcr.2017.11.012

Morrisey, E.E., Hogan, B.L.M., 2010. Preparing for the first breath: genetic and cellular mechanisms in lung development. Dev. Cell 18, 8–23. doi:10.1016/j.devcel.2009.12.010

Murakami, K., Terakado, Y., Saito, K., Jomen, Y., Takeda, H., Oshima, M., Barker, N., 2021. A genome-scale CRISPR screen reveals factors regulating Wnt-dependent renewal of mouse gastric epithelial cells. Proc. Natl. Acad. Sci. U.S.A. 118. doi:10.1073/pnas.2016806118

Musa, J., Aynaud, M.-M., Mirabeau, O., Delattre, O., Grünewald, T.G., 2017. MYBL2 (B-Myb): a central regulator of cell proliferation, cell survival and differentiation involved in tumorigenesis. Cell Death Dis 8, e2895–e2895. doi:10.1038/cddis.2017.244

Ng, A.H.M., Khoshakhlagh, P., Rojo Arias, J.E., Pasquini, G., Wang, K., Swiersy, A., Shipman, S.L., Appleton, E., Kiaee, K., Kohman, R.E., Vernet, A., Dysart, M., Leeper, K., Saylor, W., Huang, J.Y., Graveline, A., Taipale, J., Hill, D.E., Vidal, M., Melero-Martin, J.M., Busskamp, V., Church, G.M., 2021. A comprehensive library of human transcription factors for cell fate engineering. Nat. Biotechnol. 39, 510–519. doi:10.1038/s41587-020-0742-6

Nikolić, M.Z., Caritg, O., Jeng, Q., Johnson, J.-A., Sun, D., Howell, K.J., Brady, J.L., Laresgoiti, U., Allen, G., Butler, R., Zilbauer, M., Giangreco, A., Rawlins, E.L., 2017. Human embryonic lung epithelial tips are multipotent progenitors that can be expanded in vitro as long-term self-renewing organoids. Elife 6, e26575. doi:10.7554/eLife.26575

Nikolić, M.Z., Sun, D., Rawlins, E.L., 2018. Human lung development: recent progress and new challenges. Development 145, dev163485. doi:10.1242/dev.163485

Okubo, T., Knoepfler, P.S., Eisenman, R.N., Hogan, B.L.M., 2005. Nmyc plays an essential role during lung development as a dosage-sensitive regulator of progenitor cell proliferation and differentiation. Development 132, 1363–1374. doi:10.1242/dev.01678

Ostrin, E.J., Little, D.R., Gerner-Mauro, K.N., Sumner, E.A., Ríos-Corzo, R., Ambrosio, E., Holt, S.E., Forcioli-Conti, N., Akiyama, H., Hanash, S.M., Kimura, S., Huang, S.X.L., Chen, J., 2018. β-Catenin maintains lung epithelial progenitors after lung specification. Development 145, dev160788. doi:10.1242/dev.160788

Pacheco-Pinedo, E.C., Durham, A.C., Stewart, K.M., Goss, A.M., Lu, M.M., DeMayo, F.J., Morrisey, E.E., 2011. Wnt/β-catenin signaling accelerates mouse lung tumorigenesis by imposing an embryonic distal progenitor phenotype on lung epithelium. J. Clin. Invest. 121, 1935–1945. doi:10.1172/JCI44871

Planas-Paz, L., Sun, T., Pikiolek, M., Cochran, N.R., Bergling, S., Orsini, V., Yang, Z., Sigoillot, F., Jetzer, J., Syed, M., Neri, M., Schuierer, S., Morelli, L., Hoppe, P.S., Schwarzer, W., Cobos, C.M., Alford, J.L., Le Zhang Cuttat, R., Waldt, A., Carballido-Perrig, N., Nigsch, F., Kinzel, B., Nicholson, T.B., Yang, Y., Mao, X., Terracciano, L.M., Russ, C., Reece-Hoyes, J.S., Keller, C.G., Sailer, A.W., Bouwmeester, T., Greenbaum, L.E., Lugus, J.J., Cong, F., McAllister, G., Hoffman, G.R., Roma, G., Tchorz, J.S., 2019. YAP, but Not RSPO-LGR4/5, Signaling in Biliary Epithelial Cells Promotes a Ductular Reaction in Response to Liver Injury. Cell Stem Cell 25, 39–53.e10. doi:10.1016/j.stem.2019.04.005

Rawlins, E.L., Clark, C.P., Xue, Y., Hogan, B.L.M., 2009. The Id2+ distal tip lung epithelium contains individual multipotent embryonic progenitor cells. Development 136, 3741–3745. doi:10.1242/dev.037317

Ringel, T., Frey, N., Ringnalda, F., Janjuha, S., Cherkaoui, S., Butz, S., Srivatsa, S., Pirkl, M., Russo, G., Villiger, L., Rogler, G., Clevers, H., Beerenwinkel, N., Zamboni, N., Baubec, T., Schwank, G., 2020. Genome-Scale CRISPR Screening in Human Intestinal Organoids Identifies Drivers of TGF-β Resistance. Cell Stem Cell 26, 431–440.e8. doi:10.1016/j.stem.2020.02.007

Rockich, B.E., Hrycaj, S.M., Shih, H.P., Nagy, M.S., Ferguson, M.A.H., Kopp, J.L., Sander, M., Wellik, D.M., Spence, J.R., 2013. Sox9 plays multiple roles in the lung epithelium during branching morphogenesis. Proc. Natl. Acad. Sci. U.S.A. 110, E4456–64. doi:10.1073/pnas.1311847110

Sakornsakolpat, P., Prokopenko, D., Lamontagne, M., Reeve, N.F., Guyatt, A.L., Jackson, V.E., Shrine, N., Qiao, D., Bartz, T.M., Kim, D.K., Lee, M.K., Latourelle, J.C., Li, X., Morrow, J.D., Obeidat, M., Wyss, A.B., Bakke, P., Barr, R.G., Beaty, T.H., Belinsky, S.A., Brusselle, G.G., Crapo, J.D., de Jong, K., DeMeo, D.L., Fingerlin, T.E., Gharib, S.A., Gulsvik, A., Hall, I.P., Hokanson, J.E., Kim, W.J., Lomas, D.A., London, S.J., Meyers, D.A., O’Connor, G.T., Rennard, S.I., Schwartz, D.A., Sliwinski, P., Sparrow, D., Strachan, D.P., Tal-Singer, R., Tesfaigzi, Y., Vestbo, J., Vonk, J.M., Yim, J.-J., Zhou, X., Bossé, Y., Manichaikul, A., Lahousse, L., Silverman, E.K., Boezen, H.M., Wain, L.V., Tobin, M.D., Hobbs, B.D., Cho, M.H., SpiroMeta Consortium, International COPD Genetics Consortium, 2019. Genetic landscape of chronic obstructive pulmonary disease identifies heterogeneous cell-type and phenotype associations. Nat Genet 51, 494–505. doi:10.1038/s41588-018-0342-2

Sanjana, N.E., Shalem, O., Zhang, F., 2014. Improved vectors and genome-wide libraries for CRISPR screening. Nat. Methods 11, 783–784. doi:10.1038/nmeth.3047

Schmitt, M., Metzger, M., Gradl, D., Davidson, G., Orian-Rousseau, V., 2015. CD44 functions in Wnt signaling by regulating LRP6 localization and activation. Cell Death Differ 22, 677–689. doi:10.1038/cdd.2014.156

Schuijers, J., Clevers, H., 2012. Adult mammalian stem cells: the role of Wnt, Lgr5 and R-spondins. EMBO J. 31, 2685–2696. doi:10.1038/emboj.2012.149

Shih, H.P., Seymour, P.A., Patel, N.A., Xie, R., Wang, A., Liu, P.P., Yeo, G.W., Magnuson, M.A., Sander, M., 2015. A Gene Regulatory Network Cooperatively Controlled by Pdx1 and Sox9 Governs Lineage Allocation of Foregut Progenitor Cells. Cell Reports 13, 326– 336. doi:10.1016/j.celrep.2015.08.082

Smith, B.M., Traboulsi, H., Austin, J.H.M., Manichaikul, A., Hoffman, E.A., Bleecker, E.R., Cardoso, W.V., Cooper, C., Couper, D.J., Dashnaw, S.M., Guo, J., Han, M.K., Hansel, N.N., Hughes, E.W., Jacobs, D.R., Kanner, R.E., Kaufman, J.D., Kleerup, E., Lin, C.-L., Liu, K., Cascio Lo, C.M., Martinez, F.J., Nguyen, J.N., Prince, M.R., Rennard, S., Rich, S.S., Simon, L., Sun, Y., Watson, K.E., Woodruff, P.G., Baglole, C.J., Barr, R.G., Lung, F.T.M., investigators, S., 2018. Human airway branch variation and chronic obstructive pulmonary disease. PNAS 115, E974–E981. doi:10.1073/pnas.1715564115

Southall, T.D., Gold, K.S., Egger, B., Davidson, C.M., Caygill, E.E., Marshall, O.J., Brand, A.H., 2013. Cell-type-specific profiling of gene expression and chromatin binding without cell isolation: assaying RNA Pol II occupancy in neural stem cells. Dev. Cell 26, 101–112. doi:10.1016/j.devcel.2013.05.020

Sun, D., Evans, L., Perrone, F., Sokleva, V., Lim, K., Rezakhani, S., Lutolf, M., Zilbauer, M., Rawlins, E.L., 2021. A functional genetic toolbox for human tissue-derived organoids. Elife 10. doi:10.7554/eLife.67886

Tian, R., Abarientos, A., Hong, J., Hashemi, S.H., Yan, R., Dräger, N., Leng, K., Nalls, M.A., Singleton, A.B., Xu, K., Faghri, F., Kampmann, M., 2021. Genome-wide CRISPRi/a screens in human neurons link lysosomal failure to ferroptosis. Nat Neurosci 24, 1020– 1034. doi:10.1038/s41593-021-00862-0

Tian, R., Gachechiladze, M.A., Ludwig, C.H., Laurie, M.T., Hong, J.Y., Nathaniel, D., Prabhu, A.V., Fernandopulle, M.S., Patel, R., Abshari, M., Ward, M.E., Kampmann, M., 2019. CRISPR Interference-Based Platform for Multimodal Genetic Screens in Human iPSC-Derived Neurons. Neuron 104, 239–255.e12. doi:10.1016/j.neuron.2019.07.014

Tosti, L., Ashmore, J., Tan, B.S.N., Carbone, B., Mistri, T.K., Wilson, V., Tomlinson, S.R., Kaji, K., 2018. Mapping transcription factor occupancy using minimal numbers of cells in vitro and in vivo. Genome Res. 28, 592–605. doi:10.1101/gr.227124.117

Wain, L.V., Shrine, N., Artigas, M.S., Erzurumluoglu, A.M., Noyvert, B., Bossini-Castillo, L., Obeidat, M., Henry, A.P., Portelli, M.A., Hall, R.J., Billington, C.K., Rimington, T.L., Fenech, A.G., John, C., Blake, T., Jackson, V.E., Allen, R.J., Prins, B.P., Understanding Society Scientific Group, Campbell, A., Porteous, D.J., Jarvelin, M.-R., Wielscher, M., James, A.L., Hui, J., Wareham, N.J., Zhao, J.H., Wilson, J.F., Joshi, P.K., Stubbe, B., Rawal, R., Schulz, H., Imboden, M., Probst-Hensch, N.M., Karrasch, S., Gieger, C., Deary, I.J., Harris, S.E., Marten, J., Rudan, I., Enroth, S., Gyllensten, U., Kerr, S.M., Polasek, O., Kähönen, M., Surakka, I., Vitart, V., Hayward, C., Lehtimäki, T., Raitakari, O.T., Evans, D.M., Henderson, A.J., Pennell, C.E., Wang, C.A., Sly, P.D., Wan, E.S., Busch, R., Hobbs, B.D., Litonjua, A.A., Sparrow, D.W., Gulsvik, A., Bakke, P.S., Crapo, J.D., Beaty, T.H., Hansel, N.N., Mathias, R.A., Ruczinski, I., Barnes, K.C., Bossé, Y., Joubert, P., van den Berge, M., Brandsma, C.-A., Paré, P.D., Sin, D.D., Nickle, D.C., Hao, K., Gottesman, O., Dewey, F.E., Bruse, S.E., Carey, D.J., Kirchner, H.L., Geisinger-Regeneron DiscovEHR Collaboration, Jonsson, S., Thorleifsson, G., Jonsdottir, I., Gislason, T., Stefansson, K., Schurmann, C., Nadkarni, G., Bottinger, E.P., Loos, R.J.F., Walters, R.G., Chen, Z., Millwood, I.Y., Vaucher, J., Kurmi, O.P., Li, L., Hansell, A.L., Brightling, C., Zeggini, E., Cho, M.H., Silverman, E.K., Sayers, I., Trynka, G., Morris, A.P., Strachan, D.P., Hall, I.P., Tobin, M.D., 2017. Genome-wide association analyses for lung function and chronic obstructive pulmonary disease identify new loci and potential druggable targets. Nat Genet 49, 416–425. doi:10.1038/ng.3787

Weaver, M., Batts, L., Hogan, B.L.M., 2003. Tissue interactions pattern the mesenchyme of the embryonic mouse lung. Developmental Biology 258, 169–184. doi:10.1016/s0012-1606(03)00117-9

Zhang, Y., Goss, A.M., Cohen, E.D., Kadzik, R., Lepore, J.J., Muthukumaraswamy, K., Yang, J., DeMayo, F.J., Whitsett, J.A., Parmacek, M.S., Morrisey, E.E., 2008. A Gata6-Wnt pathway required for epithelial stem cell development and airway regeneration. Nat Genet 40, 862–870. doi:10.1038/ng.157

Zhang, Z., Verheyden, J.M., Hassell, J.A., Sun, X., 2009. FGF-regulated Etv genes are essential for repressing Shh expression in mouse limb buds. Dev. Cell 16, 607–613. doi:10.1016/j.devcel.2009.02.008

## Reference

Choi, H.M.T., Schwarzkopf, M., Fornace, M.E., Acharya, A., Artavanis, G., Stegmaier, J., Cunha, A., Pierce, N.A., 2018. Third-generation in situ hybridization chain reaction: multiplexed, quantitative, sensitive, versatile, robust. Development 145, dev165753. doi:10.1242/dev.165753

Li, W., Xu, H., Xiao, T., Cong, L., Love, M.I., Zhang, F., Irizarry, R.A., Liu, J.S., Brown, M., Liu, X.S., 2014. MAGeCK enables robust identification of essential genes from genome-scale CRISPR/Cas9 knockout screens. Genome Biol. 15, 554–12. doi:10.1186/s13059-014-0554-4

